# Rooting for water: Bridging Plant Hydraulics and Ecohydrology to Predict Drought Stress

**DOI:** 10.64898/2026.09.25.753735

**Authors:** Arsène Druel, Julien Ruffault, Hervé Cochard, Miquel De Cáceres, Nicolas Delpierre, Jean-Marc Limousin, Maurizio Mencuccini, Matthias Cuntz, Laurine Chir, Rémi Lemaire-Patin, Jerome Ogee, Renaud Decarsin, Sylvain Delzon, Claude Doussan, Joannès Guillemot, Emilie Joetzjer, Larter Maximilian, François Pimont, Albert Olioso, Guillaume Simioni, Léa Veuillen, Gregor Rickert, Ilhan Özgen-Xian, Nicolas K. Martin-StPaul

**Author notes:** Corresponding author: Arsène Druel (INRAE - URFM, Site Agroparc, Domaine St Paul, F - 84914 AVIGNON Cedex 9, FRANCE), **Email:**.

## Abstract

Forests regulate climate and sustain biodiversity, but their resilience is increasingly threatened by drought-induced hydraulic failure, when xylem water transport is impaired by embolism. A critical but poorly constrained determinant of this risk largely due to limited observations of deep root water uptake is the amount of root-accessible (sub)soil water storage (*S_R_*), which governs land-atmosphere exchanges during prolonged dry periods. While ecohydrological theory has long suggested a balance between soil water storage in the rooting zone, vegetation growth and drought tolerance, this concept has not yet been integrated into predictive models of hydraulic failure.

Here we propose a novel, process-based inversion framework that integrates ecohydrological optimality theory with principles of plant hydraulic to infer *S_R_*. Using the SurEau plant hydraulic model, we estimate the *S_R_* value that balances the costs of soil exploration with the avoidance of drought-induced hydraulic damage. Applied first to a well-instrumented Mediterranean *Quercus ilex* forest, the inferred *S_R_*closely matched independent neutron probe and eddy covariance measurements and accurately reproduced observed drought responses, including leaf water potential and sap flow dynamics.

We then scaled the approach across European forests remotely-sensed data, to assess spatially explicit *S_R_* estimates and associated hydraulic failure risk. Across more than 20 species and sites, inferred *S_R_* and drought stress metrics aligned with field observations and outperformed estimates based on conventional soil databases or remote-sensing-only approaches.

By linking plant physiology and ecohydrological theory within a scalable inversion framework, this approach improves predictions of forest drought risk under climate change.

**Significance Statement:** Forests rely on subsurface water to maintain functions during drought, but estimating the amount of soil water available to plants (*S_R_*) remains a major challenge. Current field-based methods cannot capture deep rooting water extraction or scale to landscape level, limiting predictions of drought impact. We present a novel, process-based approach that infers *S_R_* by combining ecohydrological optimum theory with plant hydraulic principles. We hypothesize that plants optimize rooting to balance exploration costs with the risk of hydraulic failure. Our approach predicts *S_R_* at European level, outperforming the predictions made using global soil databases or remotely-sensed estimates. This scalable framework advances understanding of forest water use and vulnerability, with broad implications for ecosystem modeling under climate change.

## Introduction

Forest trees are increasingly exposed to intensifying heat and drought worldwide, leading to severe water stress and tree mortality (Hammond et al., 2022; Sanchez-Martinez et al., 2023). By altering key ecosystem functions such as carbon uptake, transpiration and growth and disturbance regimes, including wildfires and insect outbreaks, these changes reshape forest dynamics (McDowell et al., 2020), generating feedback to the climate system (Anderegg et al., 2013). Together these impacts highlight the need to improve our ability to predict forest responses to water scarcity. Decades of ecophysiological research have established that the loss of hydraulic conductance caused by xylem embolism, that is hydraulic failure, is a primary mechanism underlying drought stress mortality risk during extreme events (Choat et al., 2012; Adams et al., 2017; Choat et al., 2018; Schuldt et al., 2020; Sanchez-Martinez et al., 2023). Plant traits governing water loss, such as leaf area, stomatal regulation, and residual conductance, together with vulnerability to xylem embolism, jointly shape both exposure to and susceptibility to hydraulic failure (Martin-StPaul et al., 2017; Choat et al., 2018; Brodribb et al., 2020). However, the occurrence of hydraulic failure at a given site emerges from the interplay among multiple factors, including climate variability, topography, soil properties, forest structure, and species traits, which has so far strongly limited our ability to predict tree and forest vulnerability to drought (Venturas et al., 2021; Trugman et al., 2020, 2021). As a result, predicting tree water stress during drought remains highly challenging (Venturas et al., 2021; Trugman et al., 2020, 2021; Choat et al., 2018; Brodribb et al., 2020).

Predicting forest water stress and the onset of hydraulic failure mainly depends on understanding how fast trees deplete the water available to them. A central component of this budget is the soil available water capacity accessible to roots (*S_R_*) (Stocker et al., 2022). *S_R_* determines the quantity of water that can be lost through transpiration thereby governing the duration trees can withstand drought before key physiological functions are impaired and hydraulic dysfunction occurs (Cochard et al., 2021; Ruffault et al., 2022, 2023). Being shaped by soil texture, stoniness, bedrock exposure, and rooting depth, the use of *S_R_*integrates the influence of aboveground drivers (rainfall and evaporative demand) and canopy condition with belowground constraints on storage and access (Gao et al., 2014). Despite its importance for understanding and predicting tree responses to drought, the spatial variability of *S_R_*remains poorly resolved (Liu et al., 2021), primarily because it is difficult to measure *in situ*. Quantifying key determinants such as soil stone content or tree rooting depth is particularly challenging (Fan et al., 2017; Chitra-Tarak et al., 2021). More importantly, a large share of *S_R_*can lie below accessible soil horizons, within weathered or fractured bedrock or groundwater reservoirs (Carrière et al., 2020; Dawson et al., 2020; McCormick et al., 2021). Consequently, most *S_R_* estimates used in vegetation models rely on field measurements, inferred rooting depths, or extrapolations from global soil databases (e.g., Piedallu et al., 2011; Tóth et al., 2017). These approaches are highly uncertain and fail to capture deep and plastic root exploration (Cabon et al., 2018; Alkassem et al., 2022; Stocker et al., 2023).

To address these limitations, several indirect approaches have been developed to estimate *S_R_*. Rather than measuring it directly, *S_R_* can be inferred from the assumption that plant canopy and subsequent water use optimally adjust to the soil properties and prevailing hydroclimate, a concept formalized as the ecohydrological equilibrium theory (EHE, Eagleson 1982). According to EHE, ecosystems under water limitation regulate canopy density (characterized by the leaf area index, or LAI), in proportion to *S_R_* in order to maximize biomass production while avoiding mortality during severe drought events (Caylor et al., 2009; Kerkhoff et al., 2004, Hoff & Rambal 2003; Schymanski et al., 2009; Cabon et al., 2018). Several studies have reported links between canopy density and water availability (Grier & Running, 1977; Schulze et al., 1996), as well as between water availability and plant hydraulic traits (Choat et al., 2012; Lens et al., 2016). Framed as a balance between water consumption, growth and water stress, the EHE concept has also proven effective in explaining both spatial and temporal variations in vegetation patterns (Caylor et al., 2009; Hoff & Rambal, 2003; Nemani & Running, 1989), and in understanding how climate regimes (in terms of precipitation distribution and evaporative demand) or stand structure influence soil moisture dynamics and status under different stomatal regulation strategies (Parolari et al., 2014; Cabon et al., 2018). Building on this framework *S_R_* has been estimated using both modelling approaches (Hoff & Rambal 2003; Schymanski et al., 2009; Cabon et al., 2018) and remote sensing (Alkassem et al., 2022; Stocker et al., 2023). These methods estimate the cumulative amount of water that an ecosystem can lose through transpiration and evaporation (using remote sensing or modelling) before reaching a stress threshold leading to embolism. This cumulative water loss is used as an indirect proxy for *S_R_*, assuming it represents the volume of water available to vegetation. Accordingly, stress thresholds are commonly defined using arbitrary soil-moisture limits or empirical stomatal-regulation functions (Schymanski, 2009; Cabon et al., 2018; Katul et al., 2007; Stocker et al., 2023). Yet these formulations neglect the hydraulic traits and water-potential thresholds that determine the onset of hydraulic failure, even though global analyses show that trees operate near their hydraulic limits (Choat et al., 2012; Manzoni et al., 2014).

We hypothesize that, because *S_R_* is ultimately constrained by the risk of hydraulic failure, integrating ecohydrological equilibrium with plant hydraulic modelling can accurately estimate *S_R_* and help to predict spatial patterns of forest drought stress. To test this hypothesis, we developed an automated algorithm that retrieves *S_R_* by inverting the process-based plant-hydraulic model SurEau (Martin-StPaul et al., 2017; Cochard et al., 2021; Ruffault et al., 2022). SurEau is a mechanistic model of water fluxes and plant water status along the soil-plant-atmosphere continuum that explicitly represents processes occurring before and after stomatal closure, including residual transpiration, xylem embolism and hydraulic failure, and depletion of internal water stores. These processes are critical for capturing plant responses to severe drought and, ultimately, drought-induced mortality. The algorithm relies on two key interrelated parameters that reflect the two theoretical assumptions underlying our framework (Fig. 1). The first parameter, baseline hydraulic risk (BHR), reflects the level of hydraulic damage, expressed as percent loss of hydraulic conductivity, that trees typically tolerate in drought years that do not reach the severity of rare, extreme events. The second parameter, hydraulic stress return period (HSRP), quantifies the frequency of drought conditions that are so extreme that they exceed the equilibrium predicted by EHE and push trees beyond their hydraulic limits, leading to widespread functional impairment and/or mortality. It is fed by LAI data widely available from remote sensing, species specific plant hydraulic traits, that are widely available in global databases. as well as soil texture and multidecadal climate time series.

**Figure 1.**
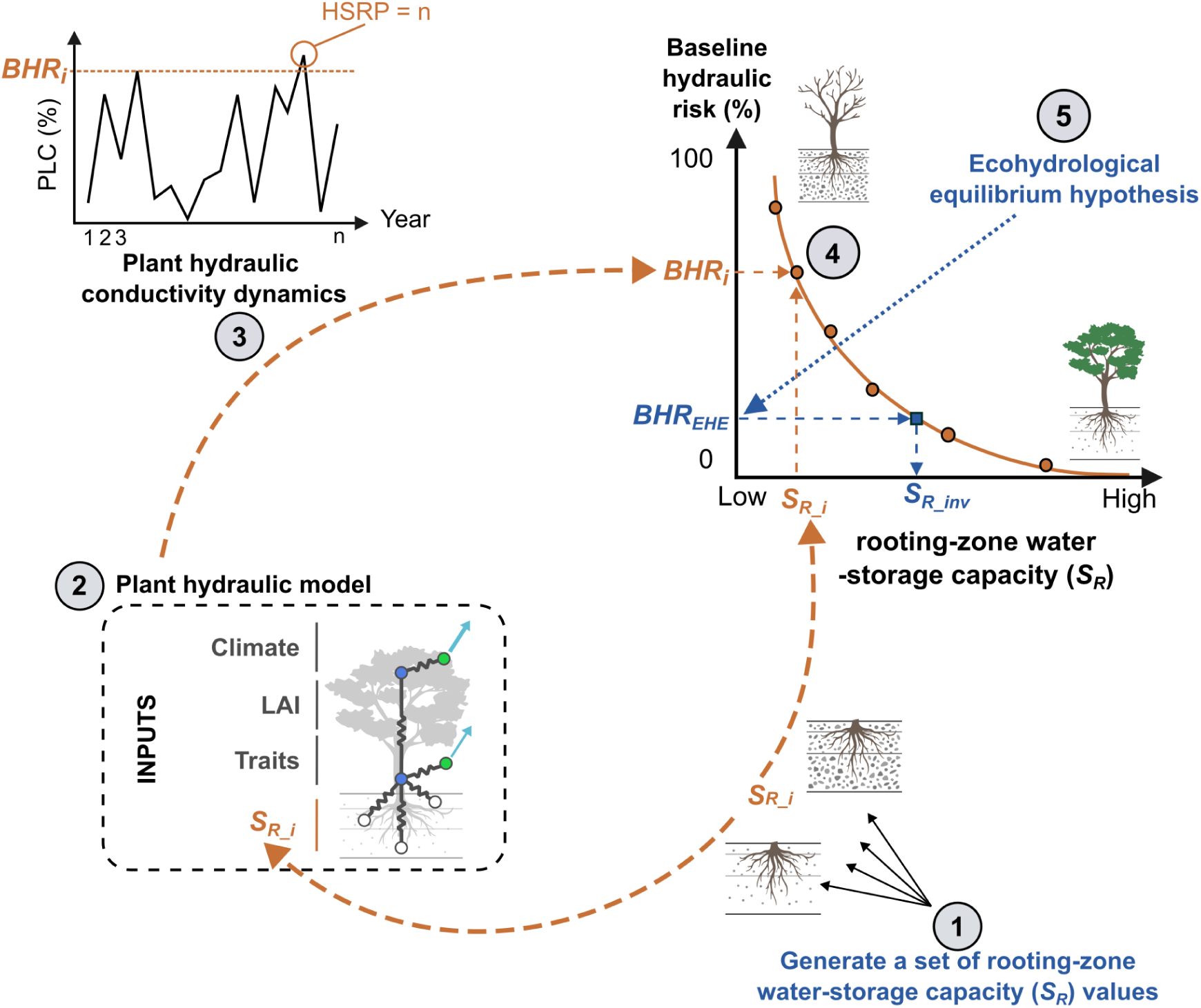
Schematic diagram of the framework applied to estimate root-zone water-storage capacity (*S_R_*) by linking ecohydrology and plant hydraulic principles. The method relies on inverting a plant hydraulic model to obtain *S_R_inv_* and involves the following steps: (1) Generate a set of potential *S_R_* values, based on a rock fraction gradient. (2) Use each *S_R_* value as a parameter of a plant hydraulic model, run under the same climate conditions over *n* years with known vegetation traits and leaf area index. (3) From model outputs, quantify the baseline hydraulic risk (*BHR_i_*) as the maximum percent loss of conductivity (PLC) reached after excluding the most extreme drought stress events. The recurrence of the most extreme years defines the hydraulic stress return period (HSRP). (4) Establish the relationship between generated *S_R_i_* values and simulated *BHR_i_*. (5) Derive *S_R_inv_* from this curve, assuming that trees follow ecohydrological equilibrium theory hypothesis and maintain baseline hydraulic risk below an acceptable threshold (*BHR_EHE_*).

We first applied the algorithm at a highly instrumented eddy-covariance site with long-term drought and sap-flow monitoring, independent *S_R_* estimates, and in situ hydraulic and stomatal traits to test the hypothesis (Ruffault et al., 2023). We then extended the approach across Europe to assess *S_R_* estimates using the ICOS database (van der Woude et al., 2023) and to evaluate predictions of extreme drought stress against a multi-species dataset of minimum water potential.

## Results

### Application of the *S_R_ inversion algorithm* at the highly instrumented site of Puechabon

We evaluated soil water storage estimates using long-term data from the Puechabon site, expecting *S_R_* to match field measurements and outperform national and European and global databases. Indeed, our framework combining EHE and hydraulic principles of the SurEau model performed well on the Puechabon site, closely matching two independent *S_R_* references derived from *in situ* data (*S_R_ref_* in Fig. 2A). First, our *S_R_* estimate (*S_R_inv_*) of 124mm +/-10 mm (Fig. 2A, Fig. S1) aligned with the value of 134 ± 2 mm, corresponding to the average maximum water consumption during drought, derived from the cumulated water deficit method (CWD, as defined in Stocker et al., 2023) applied to eddy covariance water flux measurements. Second, it was consistent with the estimate of 131 ± 6 mm derived from regular concurrent measurements of neutron probe profiles down to a 5 m depth and plant water potentials. Moreover, *S_R_inv_* departed strongly from global, European and national database values extracted for the site. According to the databases such values were either far higher (SoilGrids >220 mm) or far lower (ESDAC, IFN <50 mm) than the local estimates. Similarly, our result is much more consistent than the 260 mm estimated for the site in a recent study based on remotely sensed vegetation activity and CWD (Stocker et al., 2023).

**Figure 2.**
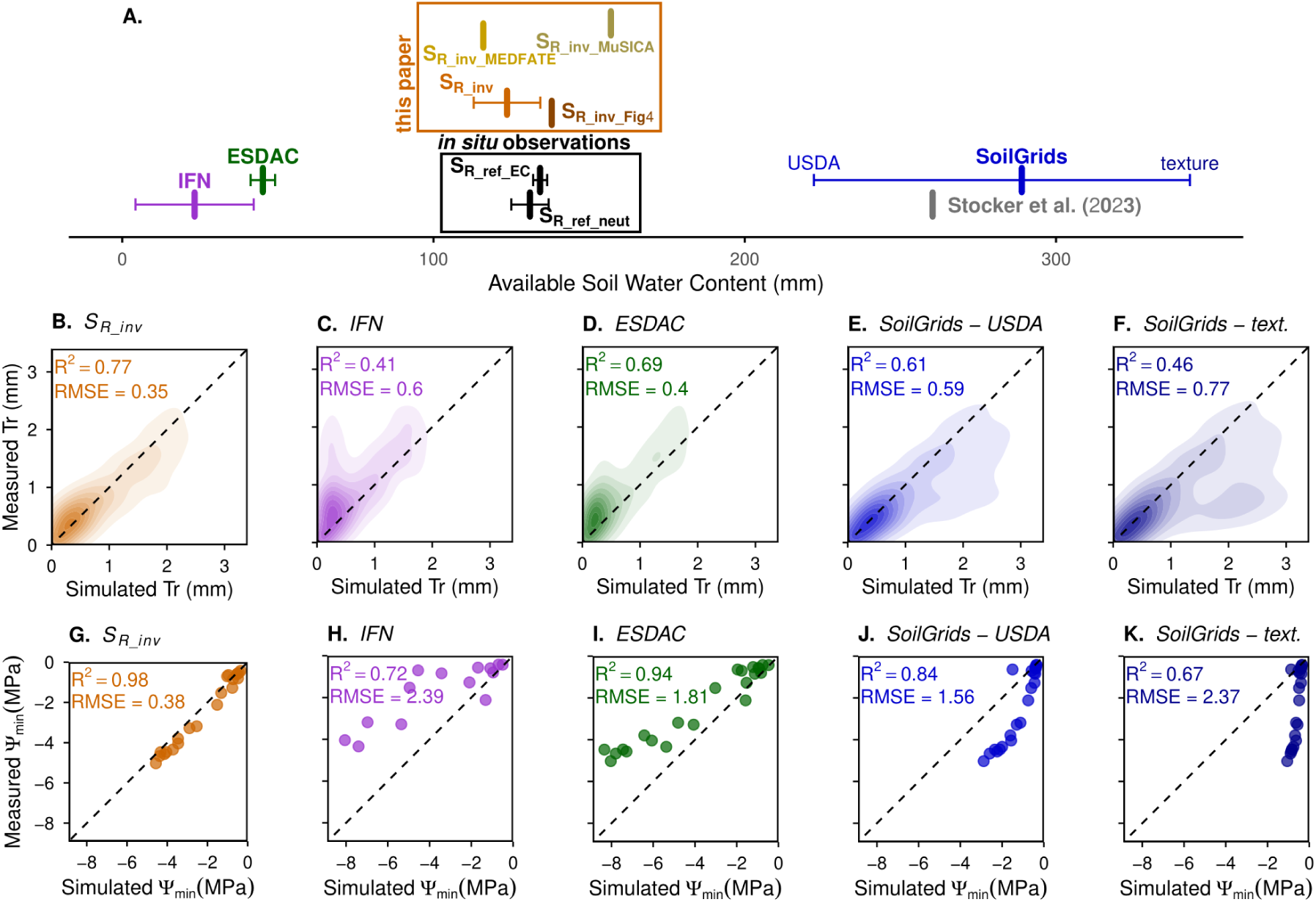
Evaluation of our ecohydrological framework grounded on plant hydraulic (presented in Fig. 1) to assess *S_R_* and drought stress at the Puechabon experimental site. Panel A compares the *S_R_inv_* derived from our *S_R_inv_ inversion algorithm* (wilting point set to −3 MPa) with *S_R_* derived from different sources. Panels B to K represent the simulation of the behaviour of trees using the different *S_R_* within a plant hydraulic model versus measured in situ on *Quercus ilex* trees from January 2016 to June 2019 on the: transpiration obtained using sapflow (panels B to F, mm) and leaf water potential *Ψ_min_* (panels G to K, MPa). In all panels, black represents the estimates from reference field data (*S_R_ref_EC_* eddy covariance and *S_R_ref_neut_* neutron probes), orange (*S_R_inv_*) the *S_R_*inversion with SurEau and Ruffault et al. (2023) configuration, brown (*S_R_inv_Fig4_*) the *S_R_* inversion using the configuration of Fig. 4A (from the application at european scale), beige (*S_R_inv_MEDFATE_* & *S_R_inv_MuSICA_*) the *S_R_* inversion with the corresponding models, purple the *S_R_*from *in situ* IFN dataset, green *S_R_* from ESDAC dataset, gray *S_R_*from map drawn up by Stocker et al. (2023) and blue *S_R_* from SoilGrids dataset (standard blue using USDA soil classes correspondence and dark blue using directly texture data, with EUPTF R package).

Using *S_R_inv_* as an input parameter in SurEau at the Puechabon site enabled accurate simulation of drought stress, including plant water potential and sap-flow dynamics (Fig. 2B–K), with performance comparable to previously published simulations in which *S_R_* was calibrated to match observed leaf water potentials (Ruffault et al 2023). This supports our implementation of the EHE framework grounded in plant hydraulic risk. Simulations made with *S_R_inv_* outperformed the simulations made with *S_R_* values from national or global databases (stronger R^2^ and lower RMSE) as well as the simulations made with the low *S_R_* (IFN and ESDAC) values extracted from local observations. Those latter strongly underestimated transpiration and overestimated drought stress (excessively low water potential; Fig. 2CDHI).

We assessed the influence of the SurEau parameters on the *S_R_inv_ inversion algorithm* by using Sobol’s methodology. We find an influential and a non-influential group of parameters (Fig. 3). The first group includes the parameter describing stand state (*LAI_max_*), as well as two species-specific traits: stomatal regulation (*Ψ_gs50_*) and vulnerability to embolism (*P_50_VC_*). Given their sensitivity values above 0.25 and exceeding the index of the “dummy parameter” (Fig. 3), these parameters are considered truly influential for the outcomes of the inversion. Among these three parameters, *LAI_max_* shows a roughly linear positive effect on *S_R_*, while *Ψ_gs50_* has a linear negative effect. In contrast, *P_50_VC_* displays a slightly increasing linear response between -9 and -7 MPa, followed by a steeper, near-exponential increase (Fig. S2). Other species traits analyzed (*K_plant,max_*, *g_res_*, *V_S_*) were not identified as significantly affecting the results of the *S_R_* inversion algorithm. Analyzing the magnitude of *S_R_inv_* variations (Table S2) induced by key SurEau parameters within the ranges observed at Puéchabon (Table 1) reveals contrasting impacts across variables (ranging from ±10 to ±25 mm), which reach up to ±35 mm when combined. Nevertheless, despite these substantial variations, the estimates remain consistent with *in situ* observations, in contrast to other databases (Fig. 2). Furthermore, assessing the impact of these *S_R_inv_* variations on tree behavior, specifically regarding transpiration and leaf water potential, shows that overall behavior remains largely unaffected (e.g., R^^2^ variations do not exceed 0.03 across studied outputs). This indicates that the acceptable parameter-induced variations in *S_R_inv_* are compensated during model simulations by the parameters themselves.

**Figure 3.**
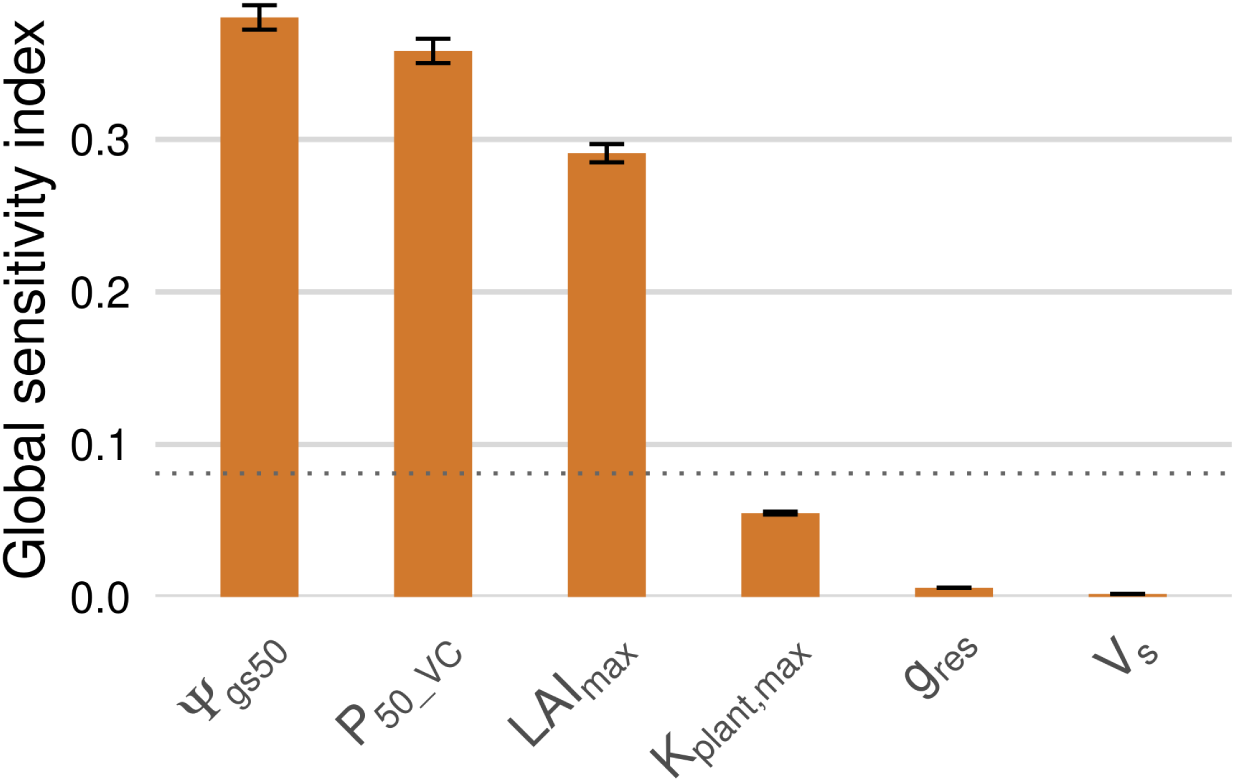
Global sensitivity index of *S_R_inv_* to plant hydraulic parameters of the model SurEau at the Puéchabon experimental site. Values shown are total-order indices with the associated standard deviation from a second-order Sobol sensitivity analysis with a 30% trait variation (see Table 1 for parameter descriptions and initial values). The horizontal dotted line indicates the estimated numerical approximation error (“dummy parameter”, value 0.081); parameters above this threshold are considered truly influential.

**Table 1.** List of parameters used in the Sobol sensitivity analysis of SurEau parameters at the Puéchabon ICOS site (FR-Pue) that may significantly influence the inversion results based on previous sensitivity analysis. Initial values derived from Ruffault et al. (2023) are also shown with the standard deviation observed on site (with associated references).

| Symbol | Variable | Unit | Value | Sd | Sd reference |
| --- | --- | --- | --- | --- | --- |
| $LAI_{max}$ | Maximum leaf area index | $m_{leaf}^2 m_{soil}^{-2}$ | 2.2 | 0.14 | Personal data |
| $\psi_{gs50}$ | Water potential causing 50% stomatal closure | MPa | -2.6 | 0.4 | Moreno et al., 2024 |
| $g_{res}$ | Residual leaf conductance water vapor at 20°C | $mmol m^{-2} s^{-1}$ | 2 | 0.3 | Limousin et al., 2022 |
| $K_{plant,max}$ | Maximum plant hydraulic conductance (per unit leaf area) from the root to the leaf | $mmol m^{-2} s^{-1} MPa$ | 0.6 | - | |
| $V_s$ | Volume of water in the stem compartment (per ground area basis) | $L m^{-2}$ | 15 | 4 | Personal data |
| $P_{50\_VC}$ | Water potential causing 50% loss of leaf or trunk hydraulic conductance. | MPa | -7 | 0.9 | Moreno et al., 2024 |

Moreover, applying the inversion algorithm with other independent plant hydraulic models yielded *S_R_inv_* estimates of the same order of magnitude as those obtained with SurEau (Fig. 2A), including 116 mm with Medfate and 157 mm with MuSICA. These estimates were also substantially closer to the local estimates (*S_R_ref_*) than those obtained from external databases. Importantly, the simulation run with inferred *S_R_inv_*values remained consistent with the simulated tree dynamics in each respective model (see Fig. S3), including transpiration (R² = 0.72 and 0.62, respectively) and leaf water potential (R² = 0.88 and 0.77, respectively).

### Toward an estimation of *S_R_* across Europe

We further explored whether the *S_R_ inversion algorithm* can help predict *S_R_* and tree hydraulic failure risk at the European scale by relying on different data sources (Fig. 4). On the one hand, Figure 4A shows that obtained *S_R_inv_*were well correlated with *S_R_ICOS_* derived from the CWD method at ICOS sites. On the other hand, *S_R_* derived from global soil databases or from the Stocker method showed much weaker agreement with *S_R_ICOS_* (Fig. 4A). The slope of the *S_R_inv_* vs. *S_R_ICOS_* regression remained >1, indicating an underestimation of some low *S_R_ICOS_*values (∼100 mm; e.g. sites of Davos, Hyytiala, and Vielsalm; Table S5) and an overestimation at few sites with high *S_R_ICOS_* (∼300 mm; sites of Hesse, Hohes Holz, and Soroe). Interestingly, the multi-species site of Font-Blanche shows an estimated *S_R_ICOS_* of 229 mm, closer to the largest *S_R_inv_*values obtained from the *S_R_ inversion algorithm* (187 mm for *Quercus ilex* and 61 mm for *Pinus halepensis*, Table S5). To preclude any climate-related bias, we tested the use of alternative climate forcing data from the SAFRAN reanalysis (Vidal et al., 2010), which provides more accurate meteorological estimates over France than ERA5. The corresponding *S_R_inv_* values were of the same order of magnitude (213 mm and 72 mm, respectively). We then assessed whether the SurEau model constrained with *S_R_inv_*was able to better predict field measurements of tree hydraulic failure risk than when constrained with *S_R_* from the SoilGrids250 database (Fig. 4B). We found that SurEau constrained with *S_R_inv_* greatly improved the prediction of predawn minimum water potential (Fig. 4B).

**Figure 4.**
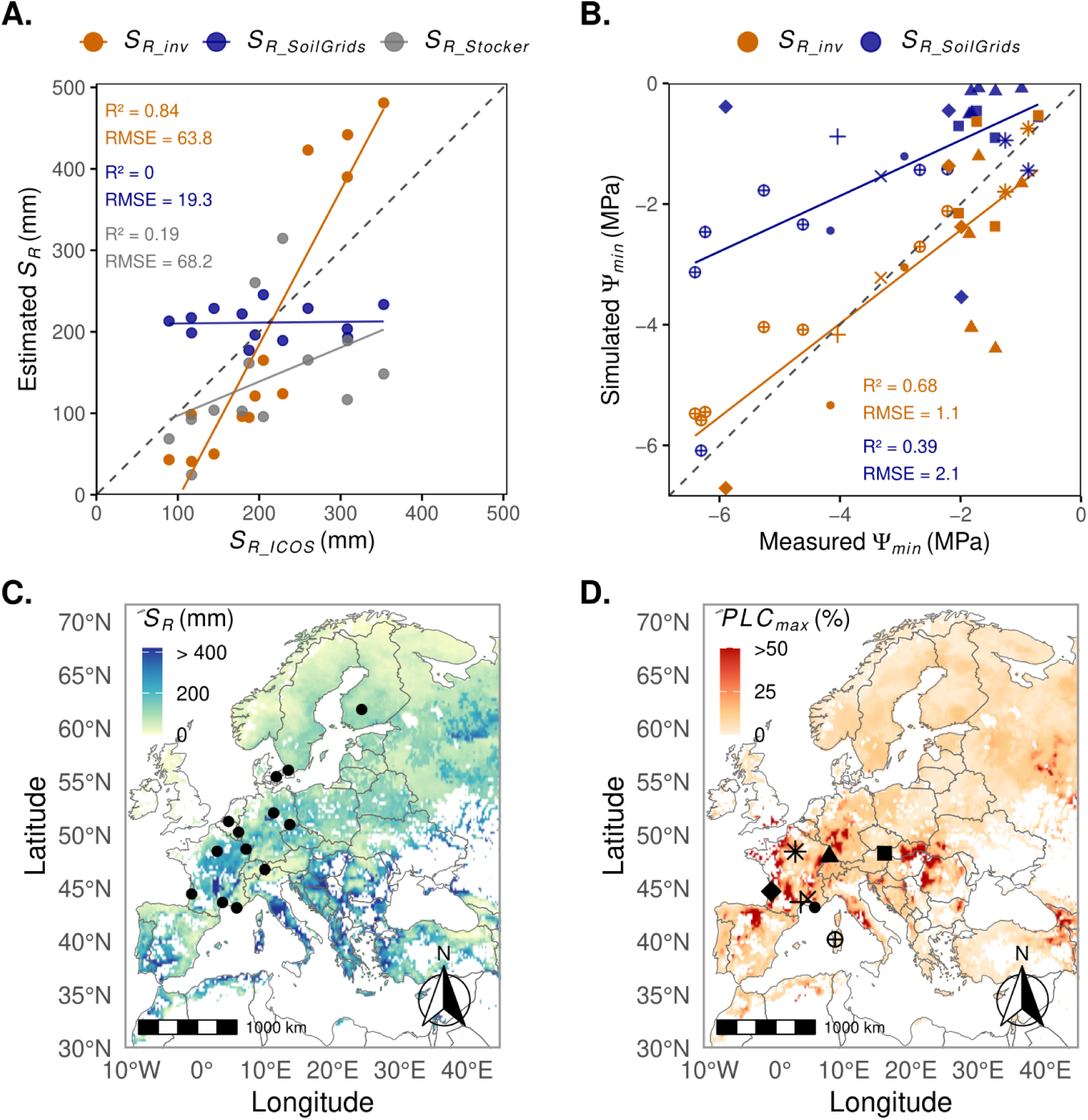
Evaluation of our ecohydrological-based estimation of SR and implications for forest drought stress across Europe. (A) Scatterplot comparing *S_R_* from the inversion algorithm (orange), SoilGrids250 (blue), and Stocker et al., 2023 (gray) against observations from thirteen ICOS sites. (B) Scatterplot of modeled drought stress (yearly minimum water potentials, *Ψ_min_*) using *S_R_* from the inversion algorithm (orange) or SoilGrids250 (blue) compared with field measurements. Species-specific and site parametrization are provided in Table S4 and S5. Maps of (C) root-accessible soil water (*S_R_*) estimated with the *S_R_*inversion algorithm at ERA5 climate resolution (0.25°) across European forests over the last three decades (the points correspond to ICOS sites; Table S5), and (D) modeled 2022 drought stress (maximum percent loss of hydraulic conductance, *PLC_max_*) based on these *S_R_* estimates using the SurEau model. In panels B and D, site symbols are: Tulln (filled square), Freiburg (filled triangle), ORPHEE (filled diamond), Macomer (circle with a cross inside), Valliguieres (cross), Barbeau (asterisk), Font-Blanche (filled circle) and Puechabon (plus), see Table S4.

To further illustrate the potential of our approach to produce estimates of *S_R_* and drought stress at large scale, we generated a European map of *S_R_inv_* (Fig. 4C) and hydraulic failure risk (Fig. 4D). Although limited by the choice of representative species, this map highlights an heterogeneous distribution of *S_R_* across Europe. Notably, boreal and high-altitude regions, such as the Alps, display particularly low *S_R_*values, consistent with observations from the northernmost ICOS site (Hyytiala, Finland) and from Davos (1689 m a.s.l.).

## Discussion

By explicitly representing tree hydraulic behaviour in an ecohydrological modelling framework we provided a mechanistic and physically consistent approach to estimating *S_R_* and drought stress. Previous large-scale estimates relied on satellite-driven crop models or water-deficit metrics that do not explicitly account for plant hydraulic functioning (Alkassem et al., 2022; Stocker et al., 2023). Our framework was successfully applied across multiple plant hydraulic models and European sites, to predict both *S_R_*and minimum water potential, a key indicator of hydraulic stress (Benito-Garzon et al., 2018; Choat et al., 2012). Its consistent performance across independent models indicates that the inversion approach is independent of the underlying hydraulic model. This evaluation across multiple sites and species supports a large-scale validity of the EHE. It also supports the assumption that hydraulic stress is an accurate optimization criterion which validates the idea that plants minimize xylem embolism under usual climatic conditions (Delzon & Cochard, 2014; Cochard & Delzon, 2013). Future studies should compare the predictions of *S_R_* and hydraulic stress using alternative optimization criteria such as growth reduction, turgor loss, hydraulic conductance reduction or stomatal regulation. Also, additional validation of the algorithm using native embolism or even direct indicators of drought damages such as leaf dieback or mortality could be used.

Importantly, our findings could pave the way for a large improvement of the spatio-temporal predictions of extreme drought stress of land vegetation, which is one of the most crucial stakes in terrestrial plant ecology as climate is rapidly heating (Hammond et al., 2022). Nowadays, most current vegetation-impacting drought assessments are based on meteorological indices or soil moisture index, that most often neglects the interacting effects of climatic aridity and other key environmental properties such as soil water stored belowground, vegetation covers, and vegetation hydraulic traits (Tramblay et al., 2020), despite they seem to be key in determining compound stress events (Li et al., 2024). The maximal belowground water stored accessible to trees (*S_R_*) has been shown to be particularly crucial as it is one of the greatest drivers of hydraulic stress in model predictions (Cochard et al., 2021; Ruffault et al., 2022; 2023). However, *S_R_* is by essence not observable at large scale. Most generally, to inform vegetation models, *S_R_*is approximated by field-based measurements of the soil water holding capacity (SWHC) derived from texture and soil depth that are interpolated (e.g. Poggio et al., 2021; Piedallu et al., 2011) sometimes using geophysical methods at local scale (Loiseau et al., 2023). Our results support that it is possible to obtain estimates of *S_R_* from aboveground data such as maximum LAI, climate and plant hydraulic traits, which are more reliable with respect to ecosystem functioning than what is provided by global soil databases that neglect rooting exploration (Fig. 2A, 4A). Our approach thereby compensates for the lack of direct belowground estimates of soil root exploration. It is complementary to recent findings supporting the prediction of *S_R_* at the plot level using only remote sensing data (Stocker et al., 2023) and EHE hypothesis (Sperry et al., 2019).

Interestingly, contrary to Stocker et al., 2023 which is built on stand-level evapotranspiration, our approach can also integrate species traits in the predictions of *S_R_*. This is illustrated by the predictions of *S_R_* at the Font-Blanche site (Fig. 4A) - an ICOS site co-dominated by trees with contrasted water use strategies (Holm oak and Aleppo pine; Moreno et al., 2021, 2024) - where we found lower values of *S_R_* for the pine (a water saver) than for the oak (a water spender) (Table S5). More importantly, once *S_R_* estimated by our algorithm was used by the SurEau model, we could tightly reproduce extreme water stress on different species and sites across Europe. Indeed, the spatial variability in predicted drought stress matches the observed patterns of forest damages (van der Woude et al., 2023). This is particularly visible in central Europe, North-Central Spain, Italy and Slovakia, where defoliation and mortality patterns correspond to the ones predicted for drought stress in this study (Hammond et al., 2022). This result demonstrates that an ecophysiologically-based approach is relevant to project future drought risk in forests, particularly by accounting for compound events, the recurrence of extreme events, and their implications for tree growth and mortality (Sohel & Marschall 2025; Torres-Ruiz et al., 2024). However, precise continent to global-scale *S_R_*estimates will require more detailed forest species maps (e.g. San-Miguel-Ayanz et al., 2016) than the simplified attribution framework used here (Supplementary S6). Such advances could further improve predictions of tree distribution and drought resilience. Importantly, our approach is not specific to SurEau but can be used with any plant hydraulic model, enabling its application with the most suitable data and modelling framework across regions.

Although our approach yields better performance in predicting *S_R_*and tree water stress than classical approaches based on model forcing using SWHC (SoilGrids) databases or multiple measurement extrapolation, several improvements are still possible, and various sources of uncertainty remain. In particular, it is sensitive to the quality of inputs. As our results suggest, climate biases (e.g. at the Font-Blanche site), which may be related to the spatial resolution of the reanalysis datasets, can strongly affect *S_R_* values and, consequently, water stress prediction. Using finer-scale data, e.g. through downscaling algorithms (De Cáceres et al., 2018; Druel et al., 2025), would likely enhance model performance (see for example the improvements with the SAFRAN data at the Font-Blanche site). Similarly, incorporating leaf area index (LAI) and species trait data at finer spatial resolutions would allow for more robust predictions.

The sensitivity analysis further showed that both LAI and species-specific hydraulic traits strongly influence the estimates of *S_R_*. Nevertheless, provided parameters remain within biologically plausible ranges, the inverted *S_R_* values retain the correct order of magnitude and perform substantially better than estimates from alternative sources. This demonstrates both a direct relationship between species-specific traits, their drought resistance strategies, and the amount of accessible soil water, as well as a strong link between vegetation state, expressed through *LAI_max_*, and *S_R_* estimation. It also highlights the strong influence of LAI measurement accuracy on the inversion results. However, the consistency between parameter values used to estimate *S_R_*and to subsequently simulate tree behavior may overlook *S_R_* uncertainties. Interestingly, the sensitivity of the *S_R_ inversion algorithm* was driven primarily by traits related to cavitation resistance and stomatal regulation: traits not identified as dominant in previous sensitivity analyses of SurEau (Cochard et al., 2021; Ruffault et al., 2022, 2023). This indicates that the inversion framework emphasizes different hydraulic processes than forward modeling alone. Moreover, in mixed-species stands where our framework is applied independently to each species, the most realistic estimate of *S_R_* appears to be the highest one. However, future developments should aim to explicitly integrate multiple species simultaneously in order to validate this assumption.

Finally, our algorithm is only valid when the EHE assumption holds - namely, when water availability is the limiting factor for maximum LAI. While many regions around the world are increasingly constrained by water stress, this assumption may not hold in areas where trees access groundwater, or in high-latitude and high-altitude regions (Fig. 4C) where light or temperature may instead be limiting factors. E.g. from a technical perspective, the algorithm may be biased in forests where LAI has recently been affected by external disturbances independent of water stress (e.g., wildfires, logging or management activities) or by abrupt changes involving long-term ecohydrological decoupling (McDowell et al., 2023). Despite this strong assumption, the use of higher-resolution data, combined with field monitoring, could help filtering out unsuitable areas and enhance the predictive potential of plant hydraulic functioning to anticipate future disturbances (Torres-Ruiz et al., 2024; Ruffault et al., 2023).

## Materials and Methods

### The “*S_R_* inversion”: an optimally based calibration method of *S_R_* based on plant hydraulic risk and EHE theory

#### General principles

The maximum amount of water that can be stored in soil and subsoil and is accessible by roots (*S_R_*) is a key parameter of vegetation models, yet this information is missing for forest ecosystems. We developed a method to assess *S_R_* from a plant hydraulic model dedicated to predict hydraulic failure under extreme drought. *S_R_* obtained from calibration (*S_R_inv_*) can be used in subsequent model runs to predict tree water stress, for example predicting plant water potential and xylem embolism. The calibration method estimates, in accordance with the ecohydrological equilibrium (EHE) theory hypotheses, the minimum required *S_R_* in order to limit the risk of reaching lethal embolism levels for a given forest stand, accounting for the specific local environmental and biological conditions (species present, climate, soil texture).

This principle was applied with the SurEau plant hydraulic model (described in the next section) and illustrated in Fig. 1. In practice, the model is configured at a given site by parametrizing (1) the forest maximum leaf area index, (2) the tree species’ most sensitive stomatal and hydraulic traits to predict the dynamic of desiccation leading to hydraulic failure (Cochard et al., 2021; Ruffault et al., 2022, see details below and Table 1), (3) the parameters of the soil moisture retention curves derived from soil texture maps, and (4) by forcing daily climate data (temperature, VPD, radiation, wind speed, rainfall) for a period representative of the normal climatic conditions of the site (in general 30 years; WMO, 2017). With such configuration, the SurEau model is applied several times, each run testing a distinct *S_R_* value (*S_R_i_)*. The different *S_R_i_*values are obtained by changing the rock fraction content over the soil profile.

To select the accurate *S_R_i_*, we defined the baseline hydraulic risk (BHR) as the maximum simulated percent loss of conductivity (PLC, %; a key metric of hydraulic failure risk) over multi-decadal periods, excluding extreme drought years based on their occurrence, quantify by the hydraulic stress return period (*HSRP*, in year; Supplementary S1). The risk level consistent with the EHE theory is referred to as *BHR_EHE_*. Indeed, for each simulation the dynamics of yearly maximum stem embolism (or PLC) was used to estimate associated *BHR_i_*defined as the maximum PLC reached under a normal climate, typically a temporal series of 30 years. Thus for each *S_R_i_* generated, a corresponding hydraulic risk (*BHR_i_*) was obtained and a relationship between *BHR_i_* and *S_R_i_*could be established. It generally resulted in a sigmoidally or a hyperbola decreasing curve. In accordance with the EHE hypothesis, which assumes that water stress is minimized during drought, a target value of baseline hydraulic risk was defined (*BHR_EHE_*) allowing us to determine the corresponding *S_R_inv_* value. The target *BHR_EHE_* should be defined to reflect hydraulic stress that does not have a substantial long-term impact on the trees, and so should represent a sufficiently low but not overly conservative hydraulic stress threshold: low enough to preserve the population, but not so high as to induce excessive sensitivity.

The influence of the two parameters required by the algorithm (HSRP and *BHR_EHE_* in Fig. 1) on *S_R_* derived from the *inversion algorithm* (*S_R_inv_*) were evaluated with SurEau configured at the Puechabon Mediterranean forest site using a Sobol sensitivity analysis (Sobol, 2001; Fig. S1). The resulting sensitivity pattern showed a steep non-linear decrease of *S_R_inv_* with the *BHR_EHE_*whereas the *HSRP* was less influential at this site. Based on ecophysiological evidence that trees operate near hydraulic limits but sustain low embolism under normal conditions (Cochard & Delzon 2013; Delzon & Cochard 2014; Arend et al., 2021) and selecting each of these two parameters within a realistic range consistent with the EHE hypothesis, we set *BHR_EHE_* to 12% and an HSRP to 10 years, keeping the uncertainty in the estimated *S_R_inv_* within ±10 mm (Supp. S1).

#### The SurEau plant hydraulic model

The SurEau model (Martin-StPaul et al., 2017; Cochard et al., 2021; Ruffault et al., 2022) was used to develop the *S_R_ inversion algorithm* in this study. SurEau is a dynamic soil-plant-atmosphere continuum model designed to simulate plant water status and fluxes within the soil-plant-atmosphere continuum during extreme drought. It has the peculiarity of representing instrumental processes that occur after the point of stomatal closure, namely transpiration losses due to residual conductance (leaf and bark), hydraulic failure due to xylem embolism, and the emptying of the plant’s internal water store, which can lead to the desiccation of plant tissue. Typical state variables include water quantity, water potential of the different compartments, and the percentage loss of hydraulic conductance between connected compartments. The model is constrained by time-varying climatic forcing (radiation, VPD, temperature, rainfall, wind speed), which are typically provided on an hourly basis. It operates on hourly to sub-hourly time steps according to stability criteria and provides outputs at hourly timesteps.

The SurEau version used here is the SurEau-Ecos published in Ruffault et al. (2022) and can be found as the *multilayersoil* branch, commit 38c4b895, at <u>forge.inrae.fr/urfm/sureau</u>. This version represents an average tree in the stand. The hydraulic architecture (Fig. 5) includes multiple soil layers and two explicit organs: a leaf and a stem. Each plant organ (i.e., leaf and stem) is described by an apoplast water reservoir (xylem and cell wall) and a symplastic compartment (living parenchyma and bark). The water volumes of these compartments are characterized by a capacitance. The connections between compartments are defined by hydraulic conductances derived from the total plant hydraulic conductance downscaled to plant organs.

**Figure 5.**
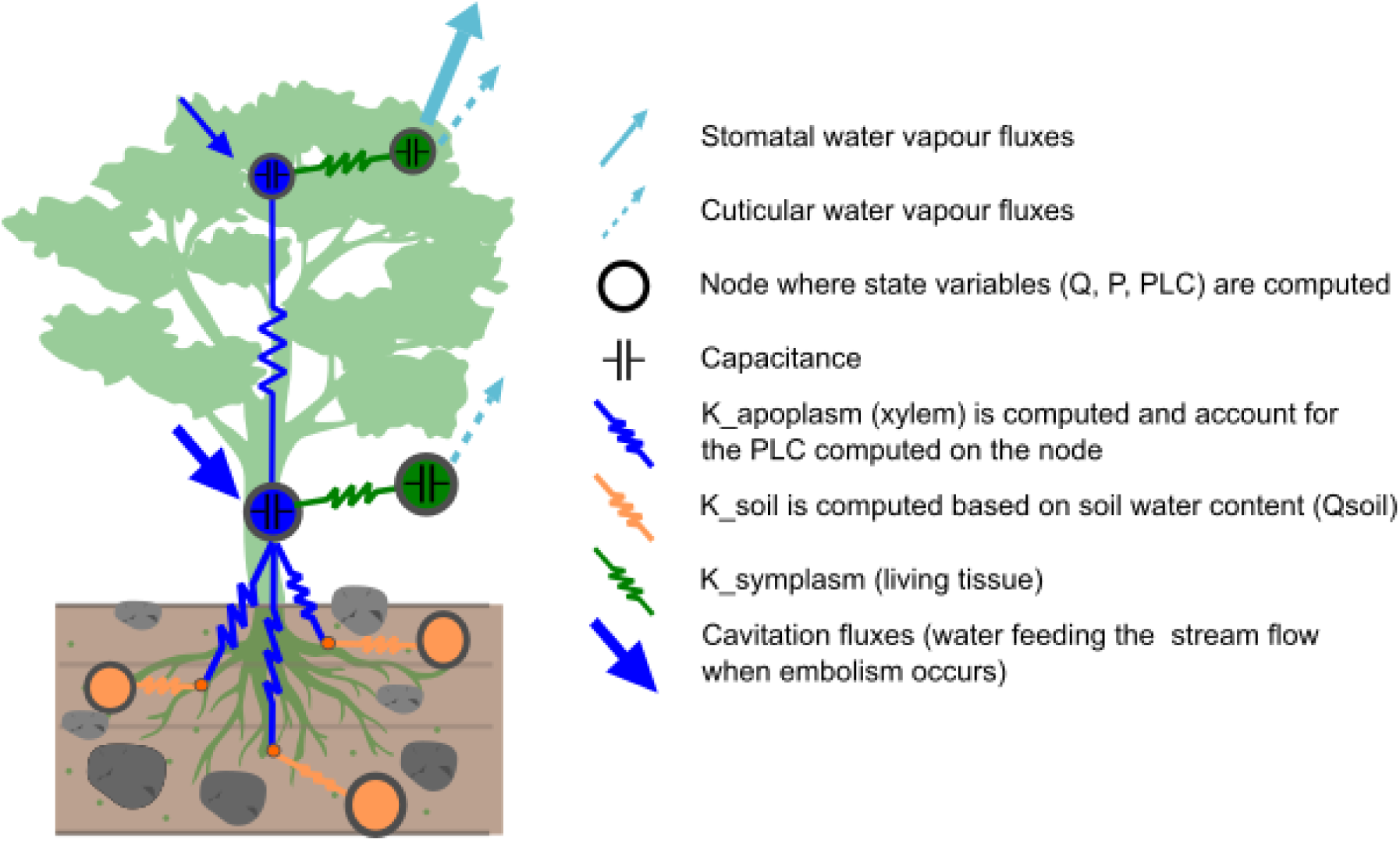
Schematic representation of the plant hydraulic architecture in SurEau-Ecos

Two different formulations can be used for the transpiration module and were used in the current study. For the application to the Puechabon site, we used the configuration of Ruffault et al. (2023) relying on the Jarvis-type stomatal model (Jarvis, 1976) for canopy-level stomatal regulation. For the application at the European scale we needed to reduce parameterization complexity of the stomatal conductance model and used the Granier-based approach (Granier et al., 1999) linking stand transpiration to potential evapotranspiration and LAI. This approach was calibrated using ICOS eddy flux series (see Supplementary S7 for calibration details).

All simulations provided in the manuscript were performed with three soil layers, except when stated differently for some sensitivity analyses (Supp. S2). The model is configured with fixed input parameters, which include soil properties and plant parameters. Soil properties include the stone fraction and the Van Genuchten parameters (e.g. Table S5), which can be defined for each layer. Plant parameters include the leaf area index, plant hydraulic traits, and parameters for the transpiration module (Table S8 and S9). Additionally, a simple one-phase phenological model is used to simulate foliage dynamics for all deciduous species. All plant hydraulic parameters that have a significant influence on hydraulic failure (defined from previous sensitivity analyses Cochard et al., 2021; Ruffault et al., 2022) were configured at the species level

### Evaluation of the *S_R_ inversion algorithm* at the Puechabon ICOS site

#### Site description

A comprehensive application of our framework is based on data collected at the Puéchabon ICOS site (FR-Pue) in the French Mediterranean region (43°44′29″N, 3°35′45″E). This forest site is dominated by *Quercus ilex L.* coppice and has remained undisturbed since the last clear-cutting event that occurred in 1942. The soil is characterized by a shallow bedrock and a high volumetric stone content of 75% in the top 0-50 cm layer and 90% below, which significantly constrains water availability (Ruffault et al., 2023; see García de Jalón et al. (2020) for more details). Meteorological data - including global radiation, temperature, precipitation, relative humidity, and wind speed required as inputs for the SurEau model - were obtained from an in-situ weather station (1999-2019 period). The site is characterized by an average temperature of 13.6°C, annual precipitation of 993 mm, and relative humidity of 67.7%. Leaf water potential (*ψ*) was measured between 2016 and 2018 at predawn and midday approximately every three weeks from May to October each year as described in Ruffault et al. (2023).

#### In situ reference S_R_ at Puechabon

We computed two different reference *S_R_* using data of eddy-covariance, soil moisture and predawn water potential from the Puechabon site (Rambal et al., 2003). First, we computed *S_R_ref_neut_*by using *in-situ* measurements of soil moisture taken using a neutron probe at six locations along a vertical profile down to 4.7 m depth at 91 different dates from 1998 to 2009. The neutron probes data were integrated along the full soil profile to provide moisture content estimate in mm. To obtain an estimate of the water content at field capacity over the profile (*WC_neut_fc_*), we used the maximal values (taken as the quantile 0.95) measured during the wet season (November to March) after removing the data measured 3 days after significant rain events (>10 mm). Then, to obtain an estimate of the water content at *Quercus ilex* wilting point (*WC_neut_wp,_* with wilting point assumed to correspond to turgor loss point) we computed the mean values (and SD) of soil moisture content for all dates with predawn water potential comprised between -2 and -4 (mean at -3.2 MPa). The *S_R_ref_neut_* estimated as the difference between *WC_neut_fc_* and *WC_neut_wp_* was 131 mm (+-6mm).

We computed an additional reference *S_R_* (*S_R_ref_EC_*) by applying the cumulated water deficit approach (CWD), based on the cumulative difference between daily-time series of eddy-covariance ETR fluxes and incoming rainfall over 15 years. We then computed the minimum water deficit values over the period as the quantile 0.1 of the minimum yearly water deficit.

#### SurEau model parameterisation

An in-depth evaluation of our framework linking ecohydrology and plant hydraulic principles to predict *S_R_* and drought stress, through the *S_R_ inversion algorithm* (see Fig. S10), was applied at the extensively monitored Puechabon site. We used the SurEau version previously applied on Quercus ilex at the Puechabon site (Ruffault et al., 2023). This version relies on a Jarvis stomatal conductance model formulation which was calibrated over sapflow measurement by adjusting two parameters of water conductances (Ruffault et al., 2023). The configuration of the model as well as the specific parametrization of other important hydraulic parameters at this species and site were retrieved from Ruffault et al. (2023), see Table S9.

#### Sensitivity analysis of S_R_inv_ to the SurEau parameters

To better understand the sensitivity of the inversion algorithm (*S_R_* inversion) to the SurEau model parameters, variance-based sensitivity analyses were performed using the Sobol method (2001). This approach allows assessing the impact of parameter variations on the results and provides ‘total-order indices’ that quantify each parameter’s contribution to output variance. The parameters were selected based on a previous study conducted under the same conditions at the same site (Ruffault et al., 2023) in order to enable comparison, and based on the identified potential impact of these parameters on the result of the inversion algorithm. The parameters selected are listed in Table 1, along with their original values, which were used to define a Sobol sample within a range of +-30%. For this purpose, the R package sensobol (version 1.1.5) was used, with the main option being a second-degree order and an initial sample size of 10k values per variable, resulting in a total of 230k simulations. The “dummy” parameter was also computed, corresponding to the estimate of the numerical approximation error (Sobol, 2001), to identify the truly influential parameters which have a higher sensitivity (see sensobol R package description).

A second round of sensitivity analysis was conducted by varying these parameters within their observed ranges at the Puechabon site (Table 1), to assess the dependence of *S_R_inv_*estimates on key model parameters and their uncertainty. Each parameter was first evaluated individually. We then jointly varied the three parameters previously identified as the strongest determinants of *S_R_inv_* in a second Sobol analysis based on 1,000 simulations, to quantify the effect of their simultaneous variation across their observed ranges on *S_R_inv_* estimates.

#### S_R_ estimates in other databases for Puechabon

Estimates of root-zone water-storage capacity (*S_R_*) are derived from various sources. The first set of data originates from rasterized databases, which provide specific values for each point. These include the SoilGrids250 database (global 2m deep-soil property maps at a spatial resolution of 250 meters based on a global compilation of soil profile data and environmental layers, Poggio et al., 2021) and the European Soil Data Centre (ESDAC) database, offering estimation at a 1,000-meter resolution derived from the European Soil Database, the Harmonized World Soil Database (HWSD), and the Soil-Terrain Database (SOTER) (Hiederer, 2013). Additionally, we obtained field measurements from the French National Forest Inventory (Inventaire Forestier National, IFN). As these local data are spatially randomized and no plot exactly matches with the study site, all plots within a 4 km radius were considered.

#### Generality of the S_R_ inversion algorithm

To assess the generality of the *S_R_ inversion algorithm* and its applicability across plant hydraulic models, we tested the approach at the same sites using two independent models, MuSICA (Ogée et al., 2003; version 3.2.8) and model (De Cáceres et al., 2023; version 5.1.1), while keeping their parameterization as consistent as possible with that used for SurEau. MEDFATE is a process-based modeling framework designed to simulate forest functioning and dynamics specially designed for Mediterranean areas. The framework allows performing soil and plant water balances in forest stands given appropriate soil, vegetation and daily weather inputs. Vegetation is represented using a set of woody plant cohorts representing individuals of the same size and species identity. MEDFATE can perform soil and plant water balances using different levels of mechanistic detail depending on submodel choices. In particular, plant hydraulics and stomatal regulation can be represented according to two different approaches: (a) steady-state plant hydraulics and optimality-based stomatal regulation (not used here); and (b) transient plant hydraulics including water compartments and empirical stomatal regulation used here (Sureau-ECOS; Ruffault et al., 2022). Compared to the original version of the ecosystem model MuSICA, several modules have been improved regarding radiative transfer, soil water transport, root water uptake or root cavitation (see McDowell et al 2016).To run MuSICA at the Puechabon site, we used a Ball-Berry model formulation for stomatal conductance (Ball et al., 1987), modified to account for stomatal closure during drought as a function of leaf water potential (Nikolov et al., 1995). Soil hydraulic properties and plant structural and functional traits were retrieved from available publications on the study site (Rambal et al, 1993; Reichstein, 2001; Limousin et al., 2010; Ruffault et al., 2023). The stomatal response to leaf water potential, as well as the leaf photosynthetic capacity and leaf cuticular conductance to water loss were then calibrated by trial and error against sap-flow and EC flux measurements over the period 2001-2013.

### European scale evaluation of the *S_R_ inversion algorithm* and application to predict drought stress

#### Global approach

A large-scale evaluation of our framework linking ecohydrology and plant hydraulic principles was performed at the European scale using multiple datasets. First, we used eddy flux time series from ICOS forest sites (van Der Woude et al., 2023) to derive independent *S_R_* estimates based on the CWD approach, which were used to validate our *S_R_inv_*estimates. Second, we built a database of *in situ* water potential measurements collected during extreme drought peaks across Europe to evaluate the ability of the SurEau model to predict associate stress once configured with *S_R_inv_*obtained from the *S_R_ inversion algorithm*. For this purpose, the SurEau plant hydraulic model was used with the Granier-type transpiration formulation. The required species-specific parameters hydraulic required were obtained from plant traits databases (Martin-StPaul et al., 2017; Larter et al., 2026) and procedures developed by Copie et al., (2025) and presented in Supplementary S8. Initial soil texture information was taken from the SoilGrids database (Poggio et al., 2021).

#### Estimation of S_R_ based on eddy-flux datasets over european ICOS forest sites

The integrated carbon observation system (ICOS) is a network composed of long-term observation stations equipped with eddy covariance systems across Europe. These stations provide high-quality, standardized data on carbon dioxide, water vapor, and energy exchanges with the atmosphere. Within this network, a consistent subset of 13 ICOS forest sites including 7 different dominant species (Table S5) were combined in the so-called Warm Winter 2020 dataset (both available on the ICOS Carbon portal), including sufficient and consistent time series bridging the datasets has been compiled (van der Woude et al., 2023). The sites represent diverse forest types, species, and climatic conditions, along a north-south gradient and adhering to standardized protocols to ensure data consistency. The data provides standardized half-hourly observations from each site, including precipitation and evapotranspiration (flux tower). This monitoring of water flux dynamics has enabled us to determine the amount of water extracted from the soil by trees via evapotranspiration. We computed the soil CWD, i.e., the cumulative difference between maximum transpired water and precipitation during extended periods without rainfall, applied to the eddy-flux data of the ICOS forest sites network. This corresponds to an estimate of the soil available water capacity functionally available for trees (*S_R_ICOS_*, Supplementary S5). The maximum of this functional *S_R_* during a period including intensive droughts, representing all water extractable by the tree from the soil, is thus considered a measured benchmark value, *S_R_ICOS_*.

#### Predicting the S_R_inv_ with our framework over european ICOS forest sites

On the same subset, we applied the *S_R_ inversion algorithm* with the SurEau model. Maximum leaf area (*LAI_max_*) were obtained from the Copernicus database (Lacaze et al., 2015); climatic data from ERA5-Land database (Muñoz-Sabater et al., 2021) from 1994 to 2023; and species from local observation (see Supplementary S4). A substantial bias was identified between local meteorological records (2016-2022) and ERA5-Land data for some sites, with precipitation at the Davos site exceeding local measurements by a factor of two, and vapor pressure deficit being underestimated by about 25% during summer at both Davos and Font-Blanche. To address this, we applied for these two sites an univariate bias correction using quantile delta mapping on ERA5-Land data (R Package MBC v0.10-7; Cannon et al., 2015). Only the dominant species at each site were considered (single species except for Tharandt and Font-Blanche, where two co-dominant species were included).

#### Predicting the drought stress over multiple sites in Europe

To assess the potential of using estimated *S_R_inv_* to predict drought stress, we evaluated the ability of the SurEau model to predict extreme drought stress across européan sites and species once SurEau was initialized with the *S_R_inv_* values. For this purpose, we built a database of field measurements of leaf water potential measured at predawn over 8 sites and 18 species over different biomes across Europe and during recent extreme drought (Moreno et al., 2024; Decarsin et al., 2024; Veuillen et al., 2026; see Table S4). These previously published measurements were conducted on experimental forest plantations or natural stands, during the peak of recent local extreme droughts (2017 or 2021 in the mediterranean or 2022 in temperate region). In situ LAI measurements previously published on these sites were used to initialize the model, using an *LAI_max_* per species where such data was available. We extracted climatic data required to run the inversion and model simulation from ERA5-Land database (Muñoz-Sabater et al., 2021), in line with *S_R_inv_*estimation over the ICOS forest sites.

#### Toward an European mapping of S_R_ and drought stress

To illustrate the potential for large-scale application of the method we have computed a spatialised estimate of *S_R_inv_* at the European level, as well as an associated drought stress assessment for the extreme drought of year 2022, through the maximum stem xylem embolism (*PLC_max_*) simulated during the year. To do this, as for the ICOS forest sites, we used the *S_R_* inversion algorithm and the SurEau model with the same configuration, forced with global data: ERA5 climate data (Muñoz-Sabater et al., 2021) from 1993 to 2022 and *LAI_max_* from the Copernicus database (Lacaze et al., 2015). As detailed in the workflow presented in Supplementary S6, in order to choose species to consider for this initial estimate, we made a rough selection of species based on the dominant forest type in each climate pixel (0.25° resolution) used as a grid for the simulations. Thus, based on the 300m vegetation distribution (ESA CCI Land Cover map; ESA, 2017) in order to extract the forest type (deciduous, broad-leaf, etc.) and based on Koppen’s climate classification (Peel et al., 2007), five major forest types and five different climate zones were defined, each associated with a species, to ultimately provide five representative but not accurate species (*Abies alba*, *Fagus sylvatica*, *Pinus pinaster*, *Quercus ilex* and *Quercus petraea*). To be consistent, given that these soil texture and *LAI_max_* databases have finer resolutions (from 1 km to 250 m) than the climate database used for these simulations (0.25° resolution), they were as well cross-referenced with the forest type indicated in the ESA CCI Land-Cover map (ESA, 2017). This makes it possible to average the LAI or textures only for grid points that correspond to tree vegetation type in the land cover map. We applied the inversion algorithm to all grid cells with more than 5% forest cover (ESA CCI Land Cover) to generate spatial estimates of *S_R_inv_*. These estimates initialized SurEau simulations for 2022 to assess drought stress from the *PLC_max_*.

## Code Availability

All software used in this study is openly available. The SR inversion algorithm is accessible at https://forge.inrae.fr/urfm/sureau_utils (files TAWinv.R and TAWinv.create.parameters.R; commit 54079244). A schematic overview of the algorithmic workflow is presented in Supplementary S10 The SurEau plant hydraulic model corresponds to version SurEau-Ecos, available at https://forge.inrae.fr/urfm/sureau (branch multilayer, commit 38c4b895).

## Acknowledgments

Druel Arsène was supported by the European Union’s Horizon 2020 Research and Innovation programme under grant agreement no. 862221 (FORGENIUS, Improving access to FORest GENetic resources Information and services for end-USers) and by the ECODIV department of INRAE (French National Research Institute for Agriculture, Food and Environment). The authors would like to thank the ICOS for providing the environmental data. We particularly thank PIs of the different forest ICOS sites studied (see Table S5). This work has received support from the French State managed by the National Research Agency under the program ANR-23-CE01-0008 TAW-Tree, the program DFG-ANR sTREssE ANR-24-CE92-0068 and France 2030 with the reference ‘ANR-24-RRII-0003’ and implemented by the INRAE EXPLOR’AE program. This work benefited from discussions conducted by the Psi-Hub working group supported by Ecodiv - INRA. We acknowledge the BioSP computing cluster at INRAE PACA for providing computational resources that supported this work.

## Author Contributions

A.D., N.M.-S., J.R., HC designed research; A.D., N.M.-S., J.R., M.D.C., N.D., J.-M.L., M.M., M.C., L.C. performed research; A.D. N.M.-S., analyzed data; R.D., J.G., S.D., E.J., L.M., G.S., L.V. provided data; A.D., N.M.-S. wrote the first draft; A.D., J.R., H.C., M.D.C., N.D., J.-M.L., M.M., M.C., L.C., R.L., J.O., R.D., S.D., C.D., J.G., E.J., L.M., F.P., A.O., G.S., L.V., G.R., I.O.-X., and N.M.-S. revised the paper.

