## Supplementary file for "Rooting for water: Bridging Plant Hydraulics and Ecohydrology to Predict Drought Stress"

##### **This PDF file includes**

Supporting text S1 to S10 (including Figures and Tables)

SI References

#### Supplementary S1: Sensitivity of the main $S_R$ inversion algorithm parameters

A Sobol (2001) sensitivity analysis of two main parameters of the $S_R$ inversion algorithm was performed in order to assess both the hydraulic risk at the baseline ecohydrological equilibrium ($BHR_{EHE}$) and so the uncertainty associated with the estimated $S_{R,inv}$. The robustness of the method was evaluated (Fig. S1) with respect to the two components of the baseline hydraulic risk (BHR) illustrated in Fig. 1. The BHR represents the maximum simulated percent loss of conductivity (PLC, %) over long periods (decades), excluding extreme drought years through the hydraulic stress return period ($HSRP$). These years correspond to rare, exceptional events that fall outside the assumptions of ecohydrological equilibrium and may result from atypical climatic conditions rather than from prevailing climatic conditions. More specifically, the $HSRP$ represents the average recurrence interval of drought events that induce substantial stem embolism (i.e. reaching a threshold embolism; Delzon & Cochard, 2014; Arend et al., 2021). The corresponding risk consistent with the ecohydrological equilibrium framework is referred to as $BHR_{EHE}$.

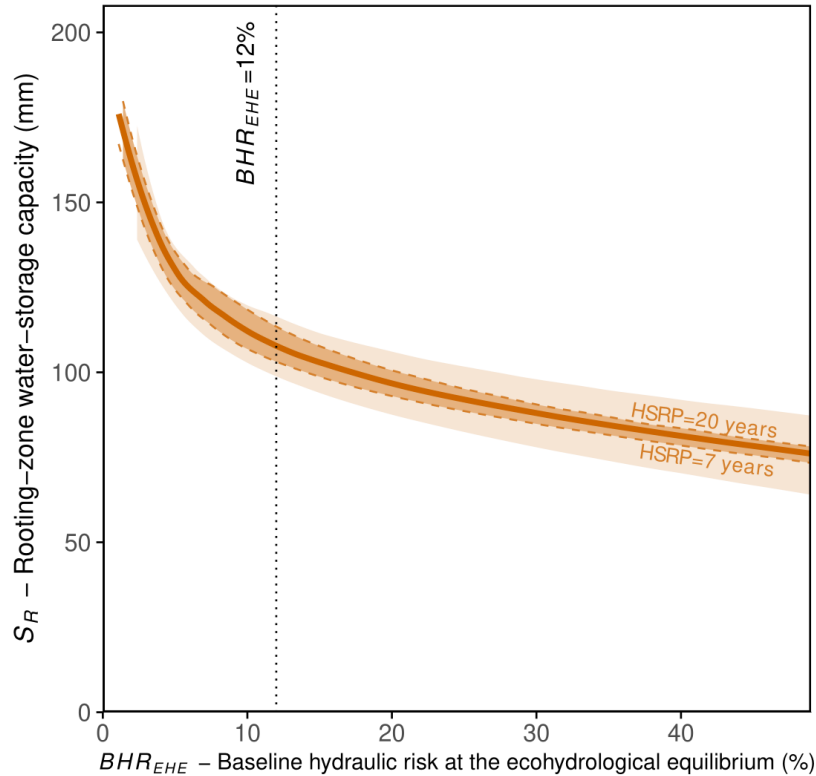

**Figure S1.** Sobol sensitivity analysis of two main parameters of the $S_R$ inversion algorithm: the baseline hydraulic stress ($BHR_{EHE}$) on the x-axis, a safety embolism threshold that does not have a substantial long-term impact on the tree population, and the hydraulic stress return period ($HSRP$) in color intensity, which reflects the acceptable frequency of extreme drought-induced hydraulic stress. In dark orange the value of $HSRP = 10$, in orange the limits of 7 and 20 years, and in light orange between 4 years (below) and without $HSRP$ (above). On a vertical black dotted line, the value of $BHR_{EHE} = 12\%$ used in the article.

For this analysis, the SurEau model configuration, plant trait parameters (species traits and LAI), and soil settings were identical to those used at the Puechabon experimental site in Fig. 2 and in Ruffault et al. (2023). Specifically, the Sobol analysis was performed using the R package sensobol (version 1.1.5), with $BHR_{EHE}$ [1-50] and $HSRP$ [4 - inf], an initial sampling of 2000 and an order defined at the 'second' level, for in total 10k simulations. The parameter ranges of the sensitivity analyses were chosen to span realistic physiological values. Overall, we observe a decreasing hyperbolic relationship between $S_R$ and $BHR_{EHE}$, while $S_R$ increases with $HSRP$. The resulting trajectories are continuous and non-erratic.

The impact of $HSRP$ depends on the $BHR_{EHE}$ range values, with different behaviour for low values ($< 5\%$), where it can affect the $S_R$ by several tens of millimetres (at constant $BHR_{EHE}$). For higher $BHR_{EHE}$ values, the influence of $HSRP$ appears more limited, with $S_R$ difference values ranging from 17 mm between extreme $HSRP$ values (a very low = 4 years or high $HSRP$ = infinite), or below 12 mm with more “reasonable” $HSRP$ values (from 7 to 20 years). Nevertheless, for extreme $HSRP$ values, starting from a $BHR_{EHE}$ of 17% ($S_R$ difference = 18 mm), the differences increase to reach almost 25 mm at a $BHR_{EHE}$ of 50%. On the contrary, for values considered reasonable, there is a continuous decrease in this difference, reaching a plateau of 5 mm from a $BHR_{EHE}$ of 38%. Consequently, an $HSRP$ between 7 and 20 years appears appropriate, with an intermediate value of 10 being reasonable, resulting in a maximum margin of error of less than 10 mm.

Moreover, the $S_R$ slope with respect to $BHR_{EHE}$ is steep for values below 6% (around -10 mm/%) showing an instability, corresponding to an absence of very slight stress that has no long-term impact, which seems necessary to avoid. At the other end, excessively high BHR values imply a long-term impact (from one year to the next), leading to an intense stress and so a break with the EHE hypothesis. A maximum limit of around 24% appears to be fairly broad, followed by a smaller variation in $S_R$ (slope below -1 mm/%). To select an intermediate value between these two limits, we performed segmented regressions analyses with the R segmented package (version 2.1-4) over a $BHR_{EHE}$ range from 6 to 24 %, and using the $HSRP$ value of 10 years. Two distinct regression segments were distinguished with slopes of -2.8 and -1.2 mm/%, respectively, and an inflection point at 12.1% ($\pm 1\%$). We therefore retained the value of $BHR_{EHE}$ of 12%, corresponding to this inflection point and to a central $S_R$ estimate within the range obtained across $BHR_{EHE}$ values from 6 to 24% (maximum difference $\sim 15$ mm). This value represents physically a moderate level of hydraulic stress with no long-term impact and is consistent with physiological reality measured in the field showing that trees operate near hydraulic limits but sustain low embolism under normal conditions (Cochard & Delzon 2013; Delzon & Cochard 2014; Arend et al., 2021).

To evaluate the uncertainty of the results and to be able to compare it to the uncertainty incoming from each variable ($BHR_{EHE}$ and $HSRP$), we used the Sobol results due to the regular sample of values. The range of $S_R$ obtained with a $BHR_{EHE}$ [6-24%] and $HSRP$ [7-20 years] is $\pm 9.4$ mm. This therefore demonstrates the interesting and consistent robustness of the results obtained using the $S_R$ inversion algorithm.

### Supplementary S2: Sensitivity of key SurEau model parameters on $S_{R\_inv}$

Figure S2 shows the response of each variable in the Sobol sensitivity analysis presented in Fig. 3. It highlights the significant but antagonistic effects of the water potential causing 50% stomatal closure ($\psi_{gs50}$ with a decreasing effect) and the water potential causing 50% loss of leaf hydraulic conductance ($P_{50\_VC}$ with an increasing effect) on $S_{R\_inv}$, as well as the influence of the leaf area index ($LAI_{max}$).

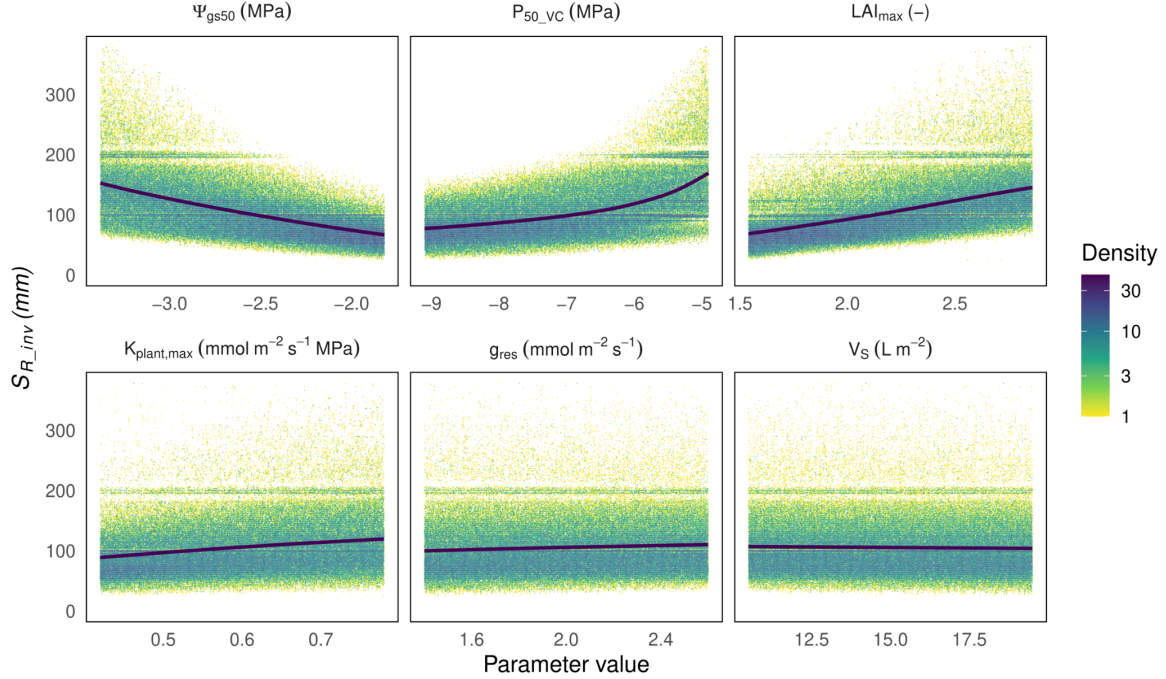

**Figure S2.** Detailed effect of key SurEau parameters on $S_{R\_inv}$ obtained with a Sobol analysis (see Fig. 3), with a wilting point set to $-1.5$ MPa. The smoothed average is shown in bold.

Below the horizontal dotted line indicating the estimated numerical approximation error (“dummy parameter”, see sensobol R package documentation), the maximum conductance from the root surface to the stem apoplasm ($K_{plant,max}$) exhibits a weak positive effect, whereas the residual conductance to leaf water vapor ($g_{res}$) and the volume of water in the stem compartment (per unit ground area, $V_s$) show no detectable influence.

The higher density of results around $S_{R\_inv} \sim 200$ mm, and to a lesser extent around 100 mm, arises from the values initially tested by the $S_R$ inversion algorithm. These initial values depend on the outcomes of previously tested $S_R$ estimates (starting with 200 mm), reflecting the adaptive search strategy of the inversion procedure.

To further investigate the effects of individual parameters on the resulting $S_{R\_inv}$ and on the functioning of the tree simulated by SurEau, we conducted a two-step sensitivity analysis based on the range of parameter variation observed at the Puechabon site (Table 1). First, we quantified the range of variation of the $S_{R\_inv}$ and the correlation with transpiration and minimum leaf water potential ($\Psi_{min}$) across the observed range of each parameter. Second, we performed a Sobol sensitivity analysis (using the same configuration as in Fig. S2, with 1,000 simulations), simultaneously varying the three parameters which have the strongest effects on $S_{R\_inv}$. This analysis assessed the sensitivity of the inversion approach to model parameters by quantifying the maximum variation in $S_{R\_inv}$ resulting from their simultaneous variation across their observed ranges. The results of these sensitivity analyses are presented in Table S2.

**Table S2.** Sensitivity analysis of key SurEau parameters on $S_{R\_inv}$ (with a wilting point set to $-1.5$ MPa) on Puechabon site and on the simulated tree response. The parameter ranges used in the sensitivity analysis are given in Table 1.

| Sensitivity analysis | $S_{R\_inv}$<br>(mm) | Transpiration (mm) | | $\Psi_{min}$ (MPa) | | $\Psi_{min}$ (MPa) | |
| --- | --- | --- | --- | --- | --- | --- | --- |
|  |  | R<sup>2</sup> | RMSE | R<sup>2</sup> | RMSE | R<sup>2</sup> | RMSE |
| Reference | 101 | 0.77 | 0.35 | 0.98 | 0.38 | 0.86 | 0.51 |
| $LAI_{max}$ only | 91 - 113 | 0.78 | 0.34-0.41 | 0.98 | 0.31-0.36 | 0.86 | 0.49-0.52 |
| $\psi_{gs50}$ only | 82 - 125 | 0.76-0.78 | 0.31-0.48 | 0.97-0.98 | 0.31-0.39 | 0.84-0.87 | 0.45-0.66 |
| $g_{res}$ only | 96 - 106 | 0.78 | 0.36-0.38 | 0.98 | 0.31-0.35 | 0.86 | 0.50-0.51 |
| $V_s$ | 99 - 103 | 0.78 | 0.37 | 0.98 | 0.32-0.34 | 0.86 | 0.50-0.51 |
| $P_{50\_VC}$ | 87 - 125 | 0.75-0.78 | 0.35-0.42 | 0.97-0.98 | 0.26-0.71 | 0.85-0.87 | 0.49-0.75 |
| Sobol with | 109 | 0.76 | 0.41 | 0.96 | 0.56 | 0.85 | 0.68 |
| $LAI_{max}$, $\psi_{gs50}$ and $P_{50\_VC}$ | $\pm 35$ | $\pm 0.03$ | $\pm 0.11$ | $\pm 0.03$ | $\pm 0.27$ | $\pm 0.03$ | $\pm 0.19$ |

Finally, sensitivity tests on the number of soil layers, total soil depth, and the distribution of rock fraction content across layers (not shown) indicated that, as long as the first two soil layers did not exceed 2 m in depth and rock fraction values remained consistent across layers (i.e., within the same order of magnitude), their effects on $S_{R\_inv}$ remained limited ($\leq 5\%$).

#### Supplementary S3:

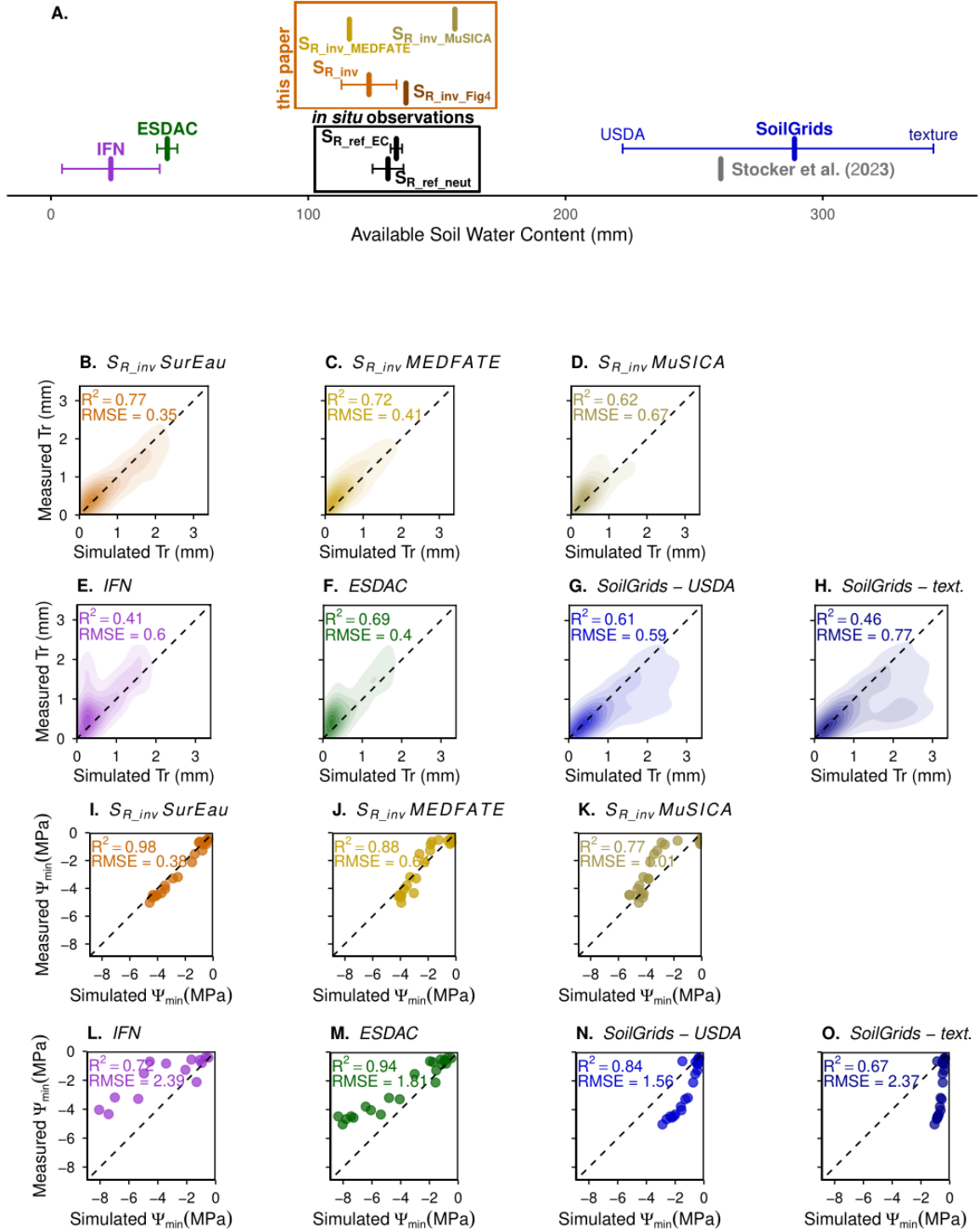

**Figure S3.** Further illustration to Figure 2, with transpiration and leaf water potential $\Psi_{min}$ simulated with the models MEDFATE and MuSICA, from respectively $S_{R\_inv\_MEDFATE}$ & $S_{R\_inv\_MuSICA}$ obtained with the $S_R$ inversion algorithm. The legend is the same as Figure 2.

**Supplementary S4:**
**European-Scale drought stress evaluation dataset**

**Table S4.** List of species and data used to apply and evaluate the $S_R$ inversion algorithm to predict drought stress at european scale (used to build Fig. 4B). The species considered, name and geographic location (longitude and latitude in degrees) of the corresponding sites, USDA soil type deduce from SoilGrids250 (Poggio et al., 2021) for each three soil layer (0-0.5, 0.5-1.5 and 1.5-4.5 m depth), maximum observed leaf area index ($LAI_{max}$) for each specie on each site (if available), measured predawn water potential during extreme drought ($\Psi_{min}$, MPa) and their published sources. The species-specific traits used to parametrize the SurEau model and apply the algorithm are presented in Supplementary S8.

| Species | Site | lon | lat | USDA soil | Date | $LAI_{max}$ obs. | $\Psi_{min}$ | $\Psi$ reference |
| --- | --- | --- | --- | --- | --- | --- | --- | --- |
| *Acer monspessulanum* | Macomer | 8.70 | 40.23 | CL;CL;CL | 2021-09-06 | 3.6 | -6.4 | Decarsin et al. (2024) |
| *Acer platanoides* | Tulln | 16.07 | 48.32 | SiL;CL;CL | 2022-07-24 | 6.0 | -1.4 | Decarsin et al. (2024) |
| *Acer platanoides* | Freiburg | 7.83 | 48.02 | L;L;L | 2022-08-10 | 3.7 | -1.8 | Decarsin et al. (2024) |
| *Acer pseudoplatanus* | Freiburg | 7.83 | 48.02 | L;L;L | 2022-08-10 | 3.7 | -1.9 | Decarsin et al. (2024) |
| *Arbutus unedo* | Macomer | 8.70 | 40.23 | CL;CL;CL | 2021-09-06 | 5.7 | -6.3 | Decarsin et al. (2024) |
| *Betula pubescens* | Freiburg | 7.83 | 48.02 | L;L;L | 2022-08-10 | 1.3 | -1.7 | Decarsin et al. (2024) |
| *Betula pendula* | Freiburg | 7.83 | 48.02 | L;L;L | 2022-08-10 | 1.3 | -1.0 | Decarsin et al. (2024) |
| *Betula pendula* | ORPHEE | -0.80 | 44.74 | SCL;SCL;SCL | 2022-08-12 | 2.1 | -2.2 | Decarsin et al. (2024) |
| *Carpinus betulus* | Barbeau | 2.78 | 48.48 | L;CL;CL | 2015-07-08 | 5.5 | -0.9 | unpublished data |
| *Carpinus betulus* | Tulln | 16.07 | 48.32 | SiL;CL;CL | 2022-07-24 | 4.9 | -1.7 | Decarsin et al. (2024) |
| *Cedrus atlantica* | Valliguières | 4.62 | 44.02 | CL;CL;CL | 2017-08-07 | 3.2 | -3.3 | Veuillen et al. (2026) |
| *Fraxinus ornus* | Macomer | 8.70 | 40.23 | CL;CL;CL | 2021-09-06 | 2.8 | -5.3 | Decarsin et al. (2024) |
| *Phillyrea latifolia* | Macomer | 8.70 | 40.23 | CL;CL;CL | 2021-09-06 | 2.6 | -6.2 | Decarsin et al. (2024) |
| *Pinus halepensis* | Font-Blanche | 5.68 | 43.24 | CL;CL;CL | 2022-08-01 | 3.1 | -2.9 | Moreno et al. (2021, 2024) |
| *Pinus halepensis* | Macomer | 8.70 | 40.23 | CL;CL;CL | 2021-09-06 | 3.0 | -2.7 | Decarsin et al. (2024) |
| *Pinus pinaster* | Macomer | 8.70 | 40.23 | CL;CL;CL | 2021-09-06 | 3.0 | -2.2 | Decarsin et al. (2024) |
| *Pinus pinaster* | ORPHEE | -0.80 | 44.74 | SCL;SCL;SCL | 2022-08-12 | 7.6 | -2.0 | Decarsin et al. (2024) |
| *Quercus ilex* | Font-Blanche | 5.68 | 43.24 | CL;CL;CL | 2022-08-01 | 3.1 | -4.2 | Moreno et al. (2021, 2024) |
| *Quercus ilex* | ORPHEE | -0.80 | 44.74 | SCL;SCL;SCL | 2022-08-12 | 1.8 | -5.9 | Decarsin et al. (2024) |
| *Quercus ilex* | Puechabon | 3.60 | 43.74 | CL;CL;CL | 2017-08-22 | 2.2 | -4.0 | Moreno et al. (2021, 2024) |
| *Quercus petraea* | Barbeau | 2.78 | 48.48 | L;CL;CL | 2016-09-21 | 5.5 | -1.3 | Postic et al. (2025) |
| *Quercus pubescens* | Macomer | 8.70 | 40.23 | CL;CL;CL | 2021-09-06 | 4.4 | -4.6 | Decarsin et al. (2024) |
| *Quercus robur* | Tulln | 16.07 | 48.32 | SiL;CL;CL | 2022-07-24 | 5.4 | -2.0 | Decarsin et al. (2024) |
| *Quercus robur* | Freiburg | 7.83 | 48.02 | L;L;L | 2022-08-10 | 1.8 | -1.4 | Decarsin et al. (2024) |
| *Quercus rubra* | Freiburg | 7.83 | 48.02 | L;L;L | 2022-08-10 | 1.8 | -1.8 | Decarsin et al. (2024) |
| *Tilia cordata* | Tulln | 16.07 | 48.32 | SiL;CL;CL | 2022-07-24 | 5.6 | -0.7 | Decarsin et al. (2024) |

The USDA full names are: C (Clay), CL (Clay Loam), L (Loam), SiC (Silt clay), SCL (Sand Clay Loam), SL (Sand Loam), SC (Sand Clay), Si (silt), LS (Loam Sand), SiCL (Silt Clay Loam), SiL (silt Loam) and S (Sand). The soil parameters of van Genuchten are directly extracted with EUPTF R package (v1.4), indicating "TOP" soil for the first soil layer.

**Supplementary S5:**
**European ICOS forest sites description**

**Table S5.** Thirteen selected ICOS forest sites (for Fig. 4A) with their dominant species, geographic location (longitude and latitude in degrees), USDA soil type deduce from SoilGrids250 (Poggio et al., 2021) for each three soil layer (0-0.5, 0.5-1.5 and 1.5-4.5 m depth), maximum leaf area index ($LAI_{max}$) from Copernicus (remote sensing; Baret et al., 2013; Lacaze et al., 2015), and soil water accessible to roots ($S_R$ in mm). Reported $S_R$ values include site-based measurements ($S_{R\_ICOS}$), inversion-based estimates ($S_{R\_inv}$), SoilGrids250-derived values ($S_{R\_SoilGrids}$), and estimates from Stocker et al. (2023) ($S_{R\_Tsocke}$).

| ICOS site | ICOS id | lon | lat | USDA soil | $LAI_{max}$ sat. | species | $S_{R\_ICOS}$ (mm) | $S_{R\_inv}$ (mm) | $S_{R\_SoilGrids}$ (mm) | $S_{R\_Tsocke}$ (mm) |
| --- | --- | --- | --- | --- | --- | --- | --- | --- | --- | --- |
| Brasschaat | BE-Bra | 4.52 | 51.31 | SL;SL;SL | 4.6 | *Pinus sylvestris* | 144 | 50 | 229 | 104 |
| Vielsalm | BE-Vie | 6.00 | 50.30 | SiL;SiL;L | 5.3 | *Fagus sylvatica* | 205 | 165 | 246 | 96 |
| Davos | CH-Dav | 9.86 | 46.81 | L;SL;SL | 3.8 | *Picea abies* | 117 | 41 | 199 | 24 |
| Hohes-Holz | DE-HoH | 11.22 | 52.09 | L;L;L | 6.2 | *Fagus sylvatica* | 353 | 480 | 233 | 148 |
| Tharandt | DE-Tha | 13.56 | 50.96 | L;L;L | 4.8 | *Picea abies*<br>*Pinus sylvestris* | 117 | 135<br>62 | 217 | 93 |
| Soroe | DK-Sor | 11.64 | 55.49 | SL;SL;SL | 6.4 | *Fagus sylvatica* | 260 | 423 | 229 | 165 |
| Hyytiala | FI-Hyy | 24.29 | 61.85 | SL;L;L | 4.5 | *Pinus sylvestris* | 89 | 43 | 213 | 69 |
| Bilos | FR-Bil | -0.96 | 44.49 | SL;SCL;SCL | 4.7 | *Pinus pinaster* | 187 | 95 | 177 | 162 |
| Font-Blanche | FR-FBn | 5.68 | 43.24 | CL;CL;CL | 2.4 | *Quercus ilex*<br>*Pinus halepensis* | 229 | 187<br>61 | 189 | 315 |
| Barbeau | FR-Fon | 2.78 | 48.48 | CL;CL;CL | 6.5 | *Quercus petraea* | 309 | 442 | 193 | 189 |
| Hesse | FR-Hes | 7.06 | 48.67 | L;CL;CL | 5.9 | *Fagus sylvatica* | 308 | 390 | 204 | 117 |
| Puechabon | FR-Pue | 3.60 | 43.74 | CL;CL;CL | 3.0 | *Quercus ilex* | 195 | 121 | 196 | 260 |
| Hyltemossa | SE-Htm | 13.43 | 56.10 | SL;SL;SL | 5.1 | *Picea abies* | 179 | 96 | 222 | 103 |

The USDA full names are: C (Clay), CL (Clay Loam), L (Loam), SiC (Silt clay), SCL (Sand Clay Loam), SL (Sand Loam), SC (Sand Clay), Si (silt), LS (Loam Sand), SiCL (Silt Clay Loam), SiL (silt Loam) and S (Sand). The soil parameters of van Genuchten are directly extracted with EUPTF R package (v1.4), indicating "TOP" soil for the first soil layer.

**Supplementary S6:**
**Continental-scale framework for scaling all inputs**
**to the climate grid for the forest $S_R$ inversion algorithm**

To produce Fig. 4C-D, we estimated the soil and subsoil water accessible to roots ($S_R$) and modeled hydraulic stress (predawn minimum water potential, $\Psi_{min}$) at the ERA5 climate resolution ($0.25^\circ$) across European forests over the last three decades, using the SurEau model with the $S_R$ inversion algorithm. In addition to ERA5 climate forcing data (1993-2022; Muñoz-Sabater et al., 2021), this required information on the maximum leaf area index ($LAI_{max}$) and soil characteristics for the dominant tree species at each grid point.

All forested areas were first extracted at 300 m resolution from the ESA CCI Land Cover map (ESA, 2017) and classified into five major forest types: broadleaved deciduous, needle-leaved evergreen, mixed forest, shrubland, and tree-shrub mosaics. Two steps followed.

(i) Selection of the tree dominant species. Dominant species, and thus associated traits, were assigned by combining the five vegetation types with Köppen bioclimatic classes (Peel et al., 2007). The latter were grouped into five broad bioclimatic zones representative of dominant European tree species (Fig. S6 & Table S6). For each vegetation-climate combination, the most frequent species was identified using the European Atlas of Forest Tree Species (San-Miguel-Ayanz et al., 2016). This approach provides a simplified, first-order representation of five dominant tree species across Europe, minimizing uncertainty in trait assignment.

(ii) Definition of $LAI_{max}$ and soil characteristics. $LAI_{max}$ was derived from Copernicus remote sensing products, at 1 km resolution before 2014 (Proba-V / SPOT-VGT) and 300 m after 2014 (Proba-V / Sentinel-3 OLCI) (Baret et al., 2013; Lacaze et al., 2015). To upscale to the $0.25^\circ$ climate grid, we selected all $LAI_{max}$ points matching each Köppen class and dominant tree type, then averaged them to obtain a representative $LAI_{max}$ per grid cell.

Soil texture was extracted from SoilGrids250 (Poggio et al., 2021) at 250 m resolution, restricted to locations where the dominant tree type occurred. Water reserves and retention curves were then computed from soil texture using the EUPTF R package (v1.4), and the dominant USDA soil texture class within each $0.25^\circ$ grid cell was retained to match the climate resolution.

Figure S6B shows the map of dominant species derived from the attribution rules defined in Table S6, restricted to grid cells where tree cover exceeded 5%.

**Table S6.** Correspondence between broad Köppen climate groups in Europe (top row), main ESA CCI Land Cover tree classes (left column), and the associated dominant tree species. The bottom row lists the Köppen class IDs included in each bioclimate group. The arid desert group (Fig. S6) corresponds to the Bwh and Bwk classes.

| **Köppen** | Xerophyllous<br>mediterranean | Temperate<br>oceanic | Central<br>continental | Boreal<br>continental | High<br>mountain |
| --- | --- | --- | --- | --- | --- |
| **ESA CCI-LC** |  |  |  |  |  |
| **Tree broadleaved**<br>**deciduous** | *Quercus petraea* | *Quercus*<br>*petraea* | *Quercus*<br>*petraea* | *Fagus*<br>*sylvatica* | *Fagus*<br>*sylvatica* |
| **Tree needleleaved**<br>**evergreen** | *Pinus pinaster* | *Pinus*<br>*pinaster* | *Abies alba* | *Abies alba* | *Abies alba* |
| **Tree mixed** | *Quercus ilex* | *Pinus*<br>*pinaster* | *Abies alba* | *Abies alba* | *Abies alba* |
| **Mosaic tree and**<br>**shrub** | *Quercus ilex* | *Quercus*<br>*petraea* | *Quercus*<br>*petraea* | *Fagus*<br>*sylvatica* | *Fagus*<br>*sylvatica* |
| **Shrubland** | *Quercus ilex* | *Quercus ilex* | *Quercus*<br>*petraea* | *Fagus*<br>*sylvatica* | *Fagus*<br>*sylvatica* |
| **Köppen ID** | Dsa, Dsb, Dsc, Bsh,<br>Bsk, Csa, Csb, Cfa | Cfb, Cfc | Dfa, Dfb | Dfc, ET, EF | ET2 |

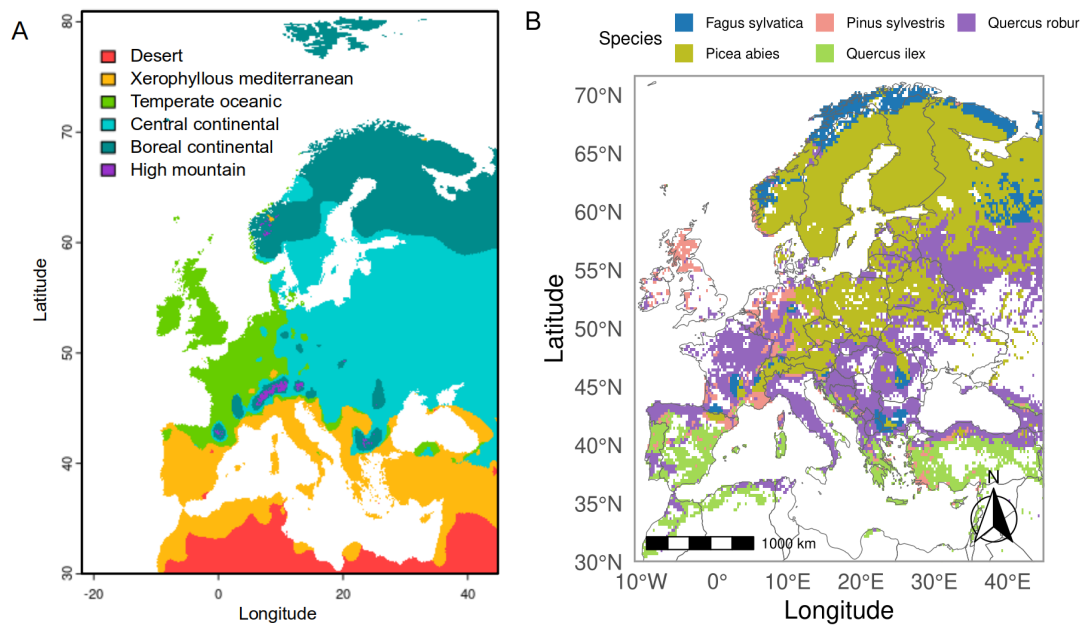

**Figure S6.** (A) Map of the Köppen bioclimatic forest groups considered in this study. See Table S6 for the Köppen classes included in each group. (B) Simplified map of dominant tree species after application of Table S6.

**Supplementary S7:**
**Procedure and data used to calibrate SurEau evapotranspiration**
**for application at european scale**

We calibrated the SurEau transpiration module (Ruffault et al 2023), at European scale by relying on the premise of Granier et al. (1999) which proposes a boundary transpiration based on Potential Evapotranspiration Transpiration and maximum LAI. It is based on the observed relationship between the ratio of the relative to potential evapotranspiration and the leaf area index (LAI). Here, we updated the parameterization of this curve using ICOS site data collected between 2016 and 2022. ICOS data are particularly suitable for this purpose because they provide harmonized measurements across multiple European sites spanning a representative north-south gradient (Fig. 4C).

To calculate the ratio $r$ under non-limiting soil water conditions, we first derived daily actual evapotranspiration (AET) from the gap-filled latent heat flux ($LE$, $W m^{-2}$):

$$AET = LE \times \frac{\Delta}{E_w \times \rho_w}$$

where $E_w$ is the evaporation heat (enthalpy) of water at 20 °C (2454 kJ kg<sup>-1</sup>), $\rho_w$ the density of water (1000 kg m<sup>-3</sup>), and $\Delta$ the time step expressed in seconds.

Daily potential evapotranspiration ($PET$) was then computed using the Priestley-Taylor formulation (Priestley and Taylor, 1972):

$$PET = \alpha \times \frac{s}{s + \gamma} \times \frac{R_n - G}{\lambda}$$

where $\alpha$ is the Priestley-Taylor coefficient (1.26), $\gamma$ the psychrometric constant (0.0666 kPa °C<sup>-1</sup>), $\lambda$ the latent heat of vaporization (2.45 MJ kg<sup>-1</sup>), $R_n$ the net radiation (MJ m<sup>-2</sup> day<sup>-1</sup>), $G$ the soil heat flux (MJ m<sup>-2</sup> day<sup>-1</sup>, assumed negligible here), and $s$ the slope of the saturation vapor pressure curve at air temperature $T$ (kPa °C<sup>-1</sup>)

With $PT_{coef}$ the Priestley-Taylor coefficient ( = 1.26), $\gamma$ the psychrometer constant ( = 0.0666 kPa °C<sup>-1</sup>), $\lambda$ the latent heat of vaporisation( = 2.45 MJ kg<sup>-1</sup>), $R_n$ the net radiation (MJ m<sup>-2</sup> day<sup>-1</sup>), $G$ the soil heat flux density (MJ m<sup>-2</sup> day<sup>-1</sup>, neglected here), et $s$ the slope of saturation vapour pressure curve at air temperature $T$ (kPa °C<sup>-1</sup>):

$$s = \frac{4098 \times 0.6108 \times \exp\left(\frac{17.27 \times T}{T + 237.3}\right)}{(T + 237.3)^2}$$

where $T$ is the mean daily air temperature (°C; Allen et al., 1998).

At each ICOS site, rainy days and the two subsequent days were excluded to minimize the influence of soil evaporation and justify neglecting $G$. To identify conditions of non-limiting soil water, we performed quantile regression (R package *quantreg* v.5.99.1) between $AET$ and $PET$ and retained the slope of the 0.8 quantile (Fig. S7A). For each site, the ratio $r$ was thus defined as this slope, and its dependence on LAI was described by a second-degree polynomial constrained through the origin (Fig. S7B). The parameters of this polynomial constrained to pass through the origin correspond to the updated Granier model values used in this study.

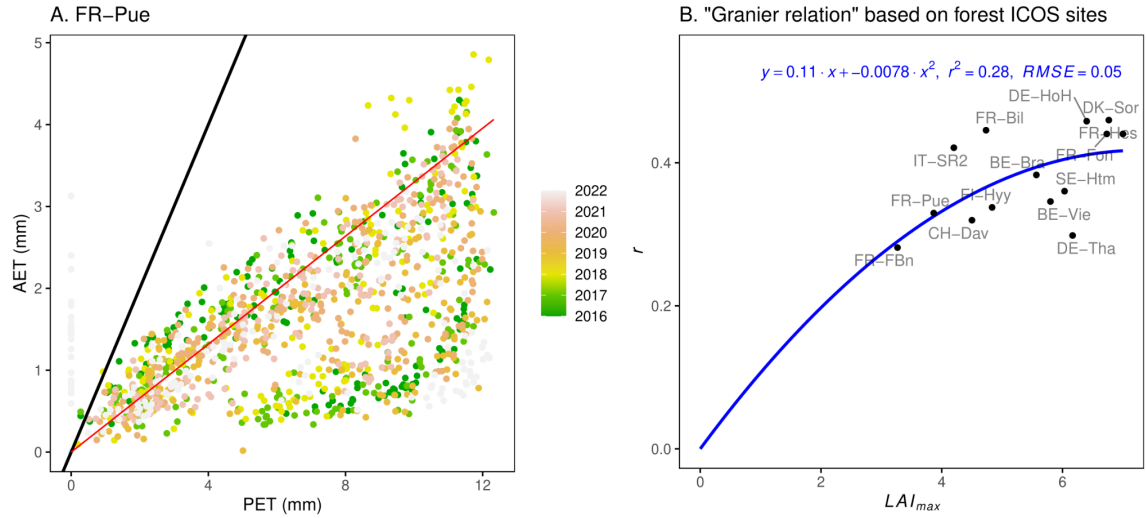

**Figure S7.** (A) Example from an ICOS site showing the relationship between relative evapotranspiration ($AET$) and potential evapotranspiration ($PET$), with the red line indicating the 0.8 quantile regression. (B) Relationship between the ratio $r$ and $LAI_{max}$ across all forest ICOS sites. In blue, the second-order polynomial regression showing the parameter values used to update the Granier model.

**Supplementary S8:**
**Species traits and parameters used to parameterize SurEau**
**for european scale application (Fig. 4)**

We parametrized SurEau for the different studied species for the European scale application only for the most sensitive traits for drought stress predictions (Table S8). We used recently published traits databases (Duursma et al., 2019 ; Burlett et al., 2025 ; Larter et al., 2026), and the procedure presented in Copie et al. (2025) to transform traits into parameters.

Specifically, the leaf residual conductance to leaf water vapor $g_{res}$ ($\text{mmol m}^{-2} \text{s}^{-1}$, per leaf projected area) was taken from Duursma et al. (2019); Billon et al. (2020) or Burlett et al. (2025). The relationship between water potential and stomatal conductance (defined by the water potential causing 88 % and 12 % of stomatal closure $\psi_{gs88}$ and $\psi_{gs12}$, MPa), as well as the maximum plant hydraulic conductance from the root surface to the leaf symplasm ($K_{plant,max}$, $\text{mmol m}^{-2} \text{s}^{-1} \text{MPa}$) were derived from the turgor loss point (TLP, MPa) taken from (Martin-StPaul et al., 2017; Kunert 2020; Larter et al., 2026), following the approach of Copie et al., 2025. The parameter of the sigmoidal vulnerability curve to cavitation (VC), defined by the water potential causing 50% loss of conductance ($P_{50,VC}$, MPa), the rate of decrease of loss of hydraulic conductance ($Slope_{VC}$, % $\text{MPa}^{-1}$). The distribution of roots along the soil profile is set as an exponential beta root profile ($\beta_{root}$; Jackson et al., 1996) and was parameterized to minimize its influence on interspecific differences in simulations, by distinguishing only between conifers ($\beta_{root} = 0.9676$) and broadleaved species ($\beta_{root} = 0.9766$).

Regarding phenology, all evergreen were assumed to have a constant LAI whereas all deciduous species were parametrized using the same single phase temperature forcing model with a sum of temperature before bud burst of 450 degree day and a leaf expansion duration of 21 days. Other parameters that have a minimal impact on drought stress are considered constant among species. They include: the leaf dry matter content (dry mass over saturated mass, $LDMC = 450 \text{ g g}^{-1}$), the leaf mass per area ($LMA$, $\text{g m}^{-2}_{leaf}$), the volume of water in the stem compartment (per ground area basis) ($V_s = 15 \text{ L m}^{-2}$), the stem and leaf apoplastic fraction ($\alpha_{Apo} = 0.4$), the stem and leaf symplasmic fraction ($\alpha_{SSym} = 0.2$ and $\alpha_{LSym} = 0.4$, respectively), the modulus of elasticity of the symplasm ($\epsilon_{Sym} = 8 \text{ MPa}$), the capacitance of the stem or leaf apoplasm ($C_{SApo} = 2.10^{-5}$ and $C_{LApo} = 1.10^{-5} \text{ mol m}^{-2}_{leaf} \text{s}^{-1} \text{MPa}^{-1}$, respectively), the canopy water storage capacity ($cws = 1.5 \text{ mm LAI}^{-1}$), the root to leaf ratio ($R_{RtoL} = 1$), the total bark (stem and branches) area to leaf ratio ($R_{TBtoL} = 0.8$), the light extinction parameter ($k = 0.5$), the leaf size ($L_s = 10 \text{ mm}$), the leaf angle ($L_a = 45^\circ$), the Priestley and Taylor coefficient ($PT_{coef} = 1.14$), the root diameter ($d_R = 2.10^{-4} \text{ m}$) and the radial conductance from the stem symplasm to the stem apoplasm ($g_{StoA} = 0.26 \text{ mmol m}^{-2} \text{s}^{-1}$).

**Table S8. Species traits parameters used for SurEau simulation in Fig. 4.**

| species | $TLP$<br>MPa | $\Psi_{gs12}$<br>MPa | $\Psi_{gs88}$<br>MPa | $K_{plant,max}$<br>$mmol\ m^{-2}\ s^{-1}$<br>MPa | $P_{50\_VC}$<br>MPa | $Slope_{VC}$<br>% $MPa^{-1}$ | $g_{res}$<br>$mmol\ m^{-2}\ s^{-1}$ |
| --- | --- | --- | --- | --- | --- | --- | --- |
| *Abies alba* | -2.70 | -1.80 | -2.70 | 1.11 | -3.71 | 97.2 | 3 |
| *Acer monspessulanum* | -3.85 | -2.57 | -3.85 | 0.78 | -6.74 | 49.6 | 4 |
| *Acer platanoides* | -1.99 | -1.33 | -1.99 | 1.51 | -4.39 | 55.5 | 4 |
| *Acer pseudoplatanus* | -2.24 | -1.49 | -2.24 | 1.34 | -2.81 | 71.9 | 4 |
| *Arbutus unedo* | -2.32 | -1.55 | -2.32 | 1.29 | -8.23 | 40.2 | 4 |
| *Betula pendula* | -1.95 | -1.30 | -1.95 | 1.54 | -2.17 | 115.1 | 4 |
| *Betula pubescens* | -1.70 | -1.13 | -1.70 | 1.76 | -1.76 | 203.9 | 4 |
| *Carpinus betulus* | -2.65 | -1.77 | -2.65 | 1.13 | -3.71 | 34.1 | 4 |
| *Cedrus atlantica* | -2.93 | -1.95 | -2.93 | 1.03 | -5.27 | 26.9 | 2.25 |
| *Fagus sylvatica* | -2.62 | -1.75 | -2.62 | 1.14 | -3.74 | 55.5 | 4 |
| *Fraxinus ornus* | -2.37 | -1.58 | -2.38 | 1.26 | -5.50 | 40.0 | 3 |
| *Larix decidua* | -2.69 | -1.79 | -2.69 | 1.11 | -3.81 | 56.6 | 3 |
| *Phillyrea latifolia* | -3.50 | -2.33 | -3.50 | 0.86 | -8.30 | 30.0 | 2.4 |
| *Picea abies* | -2.83 | -1.89 | -2.83 | 1.06 | -3.61 | 46.6 | 4 |
| *Pinus halepensis* | -2.57 | -1.71 | -2.57 | 1.17 | -4.79 | 46.7 | 1.8 |
| *Pinus pinaster* | -2.20 | -1.47 | -2.20 | 1.36 | -3.64 | 89.0 | 2.4 |
| *Pinus sylvestris* | -2.24 | -1.49 | -2.24 | 1.34 | -3.42 | 130.0 | 2.4 |
| *Pseudotsuga menziesii* | -2.97 | -1.98 | -2.97 | 1.01 | -3.48 | 76.1 | 3 |
| *Quercus ilex* | -3.15 | -2.10 | -3.15 | 0.95 | -6.63 | 30.0 | 2.4 |
| *Quercus petraea* | -2.42 | -1.61 | -2.42 | 1.24 | -4.71 | 67.0 | 3 |
| *Quercus pubescens* | -2.31 | -1.54 | -2.31 | 1.30 | -5.32 | 42.2 | 3 |
| *Quercus robur* | -2.68 | -1.78 | -2.68 | 1.12 | -4.74 | 67.1 | 4 |
| *Quercus rubra* | -2.53 | -1.69 | -2.53 | 1.19 | -4.43 | 24.5 | 4 |
| *Tilia cordata* | -2.48 | -1.66 | -2.48 | 1.21 | -3.20 | 65.2 | 3 |

**Supplementary S9:**
**Summary table of plant parametrization**

**Table S9.** List of the plant parameters, their definitions, units and indications about how they were considered in the modelling exercises (constant or variable at the individual or species level).

| Type of parameters | Parameter name | Definition | Comments on definition | Units | Set variable between species in this study | Details on parametrization |
| --- | --- | --- | --- | --- | --- | --- |
| Hydraulic parameters | EpsilonSymp_Leaf | Modulus of elasticity of the leaf symplasm | Defines Leaf symplasmic capacitance | MPa | Yes | From traits database |
|  | PiFullTurgor_Leaf | Osmotic potential at full turgor of the leaf symplasm | Defines Leaf symplasmic capacitance | MPa | Yes | From traits database |
|  | EpsilonSymp_Trunk | Modulus of elasticity of the stem symplasm | Defines Trunk symplasmic capacitance | MPa | Yes | Assumed equal to leaf (no segmentation) |
|  | PiFullTurgor_Trunk | Osmotic potential at full turgor of the stem symplasm | Defines Trunk symplasmic capacitance | MPa | Yes | Assumed equal to leaf (no segmentation) |
|  | slope_VC_Leaf | Slope of rate of leaf embolism spread at P50,L | Defines Trunk symplasmic capacitance | %.MPa<sup>-1</sup> | Yes | Assumed equal to stem (no segmentation) |
|  | P50_VC_Leaf | Water potential causing 50% loss of leaf hydraulic conductance | Leaf vulnerability curve to cavitation | MPa | Yes | assumed equal to stem (no segmentation) |
|  | slope_VC_Trunk | Slope of rate of stem embolism spread at P50,S | Leaf vulnerability curve to cavitation | %.MPa<sup>-1</sup> | Yes | From traits database |
|  | P50_VC_Trunk | Water potential causing 50% loss of stem hydraulic conductance | Leaf vulnerability curve to cavitation | MPa | Yes | From traits database |
|  | K_PlantInit | Maximum conductance from the root surface to leaf symplasm | Define the hydraulic architecture. Downscaled internally to define the hydraulic architecture | mmol.mleaf<sup>-2</sup>.s<sup>-1</sup>.MPa | NO | From traits database |
|  | K_SSymInit | Conductance from the stem apoplasm to stem symplasm | Define the hydraulic architecture | mmol.mleaf<sup>-2</sup>.s<sup>-1</sup>.MPa | NO | Not sensitive |
|  | ApoplasmicFrac_Leaf | Leaf apoplasmic fraction (from RWC leaf) |  | - | NO | From traits database |
|  | ApoplasmicFrac_Trunk | Stem apoplasmic fraction of the wood water volume |  | - | NO | Not sensitive |
|  | SymplasmicFrac_Trunk | Stem symplasmic fraction of the wood water volume |  | - | NO | Not sensitive |
|  | C_LApoInit | Capacitance of the leaf apoplasm |  | mmol.MPa<sup>-1</sup> | NO | Not sensitive |
|  | C_TApoInit | Capacitance of the stem apoplasm |  | mmol.MPa<sup>-1</sup> | NO | Not sensitive |
|  | vol_Stem | Water volume of tissue of the stem compartment (includes the root, trunk and branches) | Splitted internally between apoplasm and symplasm | L.mssoil<sup>-2</sup> | NO | Size dependant |
|  | Succulence | Leaf succulence (water content per unit leaf area) |  | g.m<sup>-2</sup> (leaf) | YES | From traits database |
|  | betaRootProfile | Shape parameter for root distribution |  | - | NO | Set high to limit sensitivity |
|  | fRootToLeaf | Root to leaf area ratio |  | - | NO | From traits database |
|  | rootRadius | Fine root diameter |  | m | NO | From traits database |
| Stomatal regulation | P50_gs | Water potential causing 50% stomatal closure |  | MPa | Yes | Derived from TLP |
|  | slope_gs | Rate of decrease in stomatal conductance at gs,50 |  | %.MPa<sup>-1</sup> | Yes | Derived from TLP |
| **(1)**<br>**Transpiration mode for the application at Puechabon : Jarvis stomatal model for (used for the Puechabon Application)** | gsMax | Maximum stomatal conductance |  | mmol.mleaf<sup>-2</sup>.s<sup>-1</sup> | NO | Calibrated using sapflow at Puechabon |
|  | JarvisPAR | Response of g<sub>stom</sub> to light |  | - | NO | Calibrated using sapflow at Puechabon |
|  | Tgs_optim | Temperature at maximal stomatal conductance |  | °C | NO | Calibrated using sapflow at Puechabon |
|  | Tgs_sens | Stomatal sensitivity to temperature |  | °C | NO | Calibrated using sapflow at Puechabon |
|  | gCrown0 | Reference crown conductance |  | mmol.mleaf<sup>-2</sup>.s<sup>-1</sup> | NO | Calibrated using sapflow at Puechabon |
| **(2)**<br>**Transpiration model for application at european scale: Granier** | Three empirical parameters of the model with relationships between Transpiration/ETP and LAI (used for transpiration at ICOS European network and at european scale) |  | - | - | NO | Calibrated using ICOS ETR data see Supplementary 5. |
| Residual and bark transpiration | gmin20 and gbark | Cuticular conductance at 20°C |  | mmol.mleaf<sup>-2</sup>.s<sup>-1</sup> | Yes | Database (gbark set equal to gmin20) |
|  | Q10_1_gmin | Temperature dependance of g<sub>min</sub> |  | - | NO | From traits database |
|  | Q10_2_gmin | Temperature dependance of g<sub>min</sub> |  | - | NO | From traits database |
|  | TPhase_gmin | Temperature for transition phase of g<sub>min</sub> |  | °C | NO | From traits database |

### Supplementary S10:

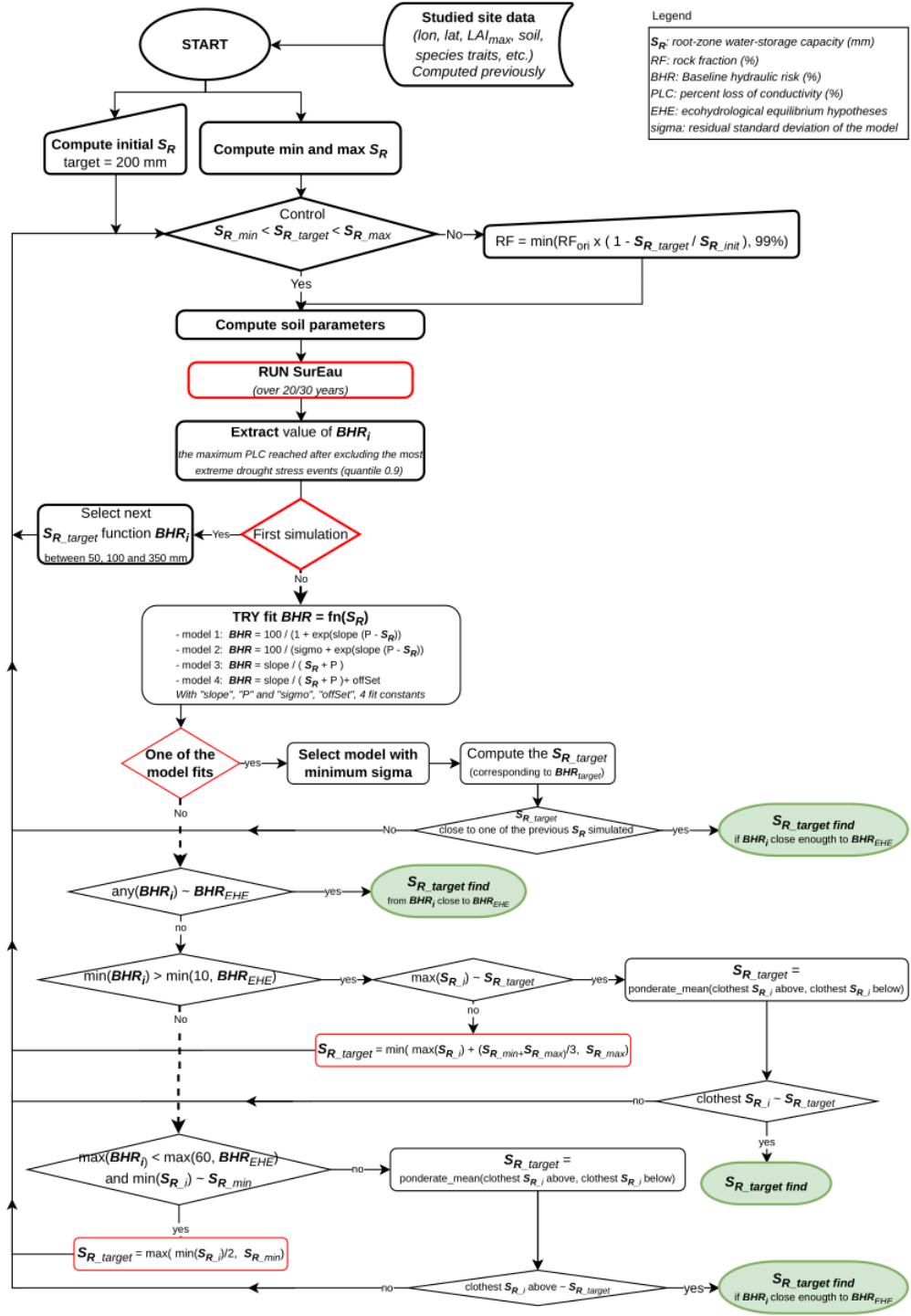

**Figure S10.** Flowchart of the $S_R$ inversion algorithm.

This figure illustrates the steps of the framework, which incorporates an optimization procedure to reduce the number of simulations required to estimate $S_{R\_inv}$ while improving precision. The algorithm begins with a fixed $S_R$ value (200 mm), then selects a second value based on the initial result. It subsequently relies on fitted functions or simple linear regressions between the closest simulation outputs to identify the $S_R$ value that satisfies the ecohydrological equilibrium hypothesis while maintaining baseline hydraulic risk below an acceptable threshold ($BHR_{EHE}$).
